# Mechanisms of mucosal immunity to oral *Shigella* infection in a physiological mouse model

**DOI:** 10.64898/2026.09.08.750193

**Authors:** Janet Peace Babirye, Emma F. Lackner, Sudyut Yuvaraj, Roberto A. Chavez, Charlotte A. Nichols, Kevin D. Eislmayr, Stefan A. Fattinger, Dmitri I. Kotov, Cammie F. Lesser, Russell E. Vance

## Abstract

*Shigella flexneri* causes bacillary dysentery, a diarrheal disease responsible for significant global morbidity and mortality. Despite extensive efforts, there is no licensed *Shigella* vaccine, and due to the lack of tractable and physiological models, mechanisms of adaptive immunity to *Shigella* are poorly understood. Here, we establish a mouse model that permits mechanistic dissection of adaptive immunity to a physiological oral challenge with *Shigella*. We find primary *Shigella* infection confers robust cross-serotype protection against secondary challenge, in a manner strictly dependent on the adaptive immune compartment. *Shigella* infection induces *Shigella*-specific CD4^+^ and CD8^+^ T cells, but only CD4^+^ T cells are required for protection. CD4^+^ T cells produce IFN*γ* upon secondary challenge, and help B cells produce *Shigella*-specific IgA. Neither anti-*Shigella* antibodies nor IFN*γ* are individually required for immunity to *Shigella*, but loss of both eliminates protective immunity. Collectively, our results demonstrate that CD4^+^ T cells orchestrate antibody and cytokine defense against shigellosis.

## Introduction

*Shigella* species cause bacillary dysentery, a severe diarrheal disease that imposes a disproportionate burden on children under five years of age in low- and middle-income countries^1–3^. *Shigella* is the second leading cause of pediatric diarrheal mortality^4^. Among survivors, *Shigella* can inflict persistent morbidity, notably stunting growth and impairing cognitive development in young children^5^. Because a licensed vaccine against *Shigella* remains unavailable despite decades of research, clinical management relies heavily on the use of antibiotics. As a result, the spread of multidrug-resistant *Shigella* strains is an emerging threat, highlighting the urgent need for an effective vaccine^6–8^.

*Shigella* transmission occurs via the fecal-oral route after ingestion of contaminated food or water^9^. The infectious dose for humans is variable but can be as low as 100 bacteria^10^. Upon reaching the intestinal lumen, the bacterium utilizes a virulence plasmid-encoded Type III Secretion System (T3SS) to translocate effector proteins into host intestinal epithelial cells (IECs), triggering bacterial internalization^11–14^. Following entry, *Shigella* escapes into the cytosol where it replicates rapidly. *Shigella* evades extracellular immune defense by spreading directly from cell to cell using a cell surface protein IcsA (VirG) that co-opts and polymerizes host actin to propel bacteria into adjacent cells^15,16^.

A major bottleneck in *Shigella* vaccine development has been the lack of a suitable animal model, as humans are the sole natural host. Epidemiological data indicate that serum IgG targeting lipopolysaccharide (LPS) O-antigen is a primary correlate of protection in humans^17–19^. Furthermore, controlled human challenge infection studies demonstrate that IgG responses to VirG and the T3SS protein IpaB strongly correlate with protection against shigellosis^20,21^. Beyond serum IgG, Phase 2b clinical trials have revealed concurrent increases in circulating *S. flexneri* LPS-specific IgA antibody-secreting cells within PBMCs and elevated serum IgA titers^22,23^. In parallel, oral immunization of human volunteers with the live-attenuated candidate CVD 1208S elicits robust phenotypic activation across circulating CD4^+^ and CD8^+^ T cell compartments^24^. However, moving beyond correlates of immunity to identify causal mechanisms of protective immunity requires tractable animal models in which specific components of host adaptive immunity can be experimentally manipulated or eliminated.

Because wild-type mice are fully resistant to oral *Shigella* infection, studies of adaptive immunity have largely relied on a broncho-pulmonary mouse model that does not recapitulate the physiological route of infection or natural site of bacterial replication^25,26^. While early studies suggested that protective immunity to *Shigella* is antibody-mediated and develops independently of T lymphocytes or interferon gamma (IFN*γ*)^27–30^, more recent work has highlighted a critical requirement for helper T cells, specifically Th17 cells, in restricting bacterial growth during re-infection^31^. Evidence from the broncho-pulmonary model also suggests that CD8^+^ T cells are not primed during primary exposure and contribute minimally to secondary protection^32^. Whether such observations hold true during infection of the gastrointestinal tract remains unknown.

We recently reported that the differential susceptibility of mice and humans to oral *Shigella* infection stems from key functional differences in the innate immune response of intestinal epithelial cells (IECs)^33,34^. Unlike humans, mice express high levels of the NAIP–NLRC4 inflammasome in IECs. This inflammasome detects the T3SS of *Shigella* and other bacteria, leading to rapid pyroptosis and expulsion of infected cells from the epithelium^35,36^. In addition to its low expression in human IECs ^37^, human NLRC4 is also inhibited by a secreted *Shigella* effector, OspF^38^. Likewise, human (but not mouse) gasdermin D, a pore-forming protein that mediates pyroptosis downstream of NAIP–NLRC4, is inhibited by a different *Shigella* effector, IpaH7.8^39^. We found that genetic ablation of the NAIP-NLRC4 inflammasome renders mice highly susceptible to oral *Shigella* infection, and leads to self-limiting pathology—including diarrhea, overt intestinal inflammation, and bacterial replication in IECs—that closely mirrors the typical disease progression seen in humans^33,34^. *Nlrc4*^−/−^ mice represent a near ideal humanized model since they recapitulate the primary reason for human susceptibility to *Shigella*, namely, a lack of functional NAIP–NLRC4 in IECs. One key difference between our oral *Shigella* model and natural human infection is that our *Nlrc4*^−/−^ mice are pretreated with streptomycin prior to oral challenge. This treatment, also commonly utilized in other oral infection mouse models^40^, is necessary to overcome the colonization resistance of the gut lumen provided by the mouse commensal microbiota. Although streptomycin helps facilitate initial colonization of the gut lumen, the subsequent events of bacterial invasion, replication, and host-mediated clearance closely mimic what is seen in humans. Our studies of the innate response to *Shigella* have established that macrophages and IFN*γ* are key players that orchestrate innate immune clearance of a primary *Shigella* infection^41^. However, whether the adaptive immune system also plays a role in facilitating immune clearance of *Shigella* in the gut remains unknown.

Here we demonstrate that a primary *Shigella* infection elicits an adaptive immune response that protects against a secondary *Shigella* challenge. We found that primary infection elicits *Shigella*-specific CD4^+^ and CD8^+^ T cells. However, protective immunity only requires CD4^+^ T cells, whereas CD8^+^ T cells are dispensable. We found that CD4^+^ T cells are required to help B cells mount a *Shigella*-specific mucosal IgA response, and in addition, directly produce IFN*γ* upon re-challenge. Surprisingly, neither B cell depletion nor IFN*γ* neutralization individually abrogated immunity to *Shigella*, suggesting that B cell-mediated humoral immunity and CD4^+^ T cell-derived IFN*γ* provide a layered defense against infection. Our results provide essential insights into the nature of protective immunity to *Shigella* that are directly relevant to ongoing efforts to develop a *Shigella* vaccine.

## Results

### An experimental system in which primary *Shigella* infection protects against re-challenge

To dissect the mechanisms of adaptive immunity to *Shigella flexneri*, we first sought to establish a physiological and experimentally tractable system in which adaptive immunity is elicited and protects against subsequent infection. We initially asked if *Nlrc4*^−/−^ mice that recovered from a primary oral *Shigella flexneri* 2457T (serotype 2a) infection, were resistant to a subsequent re-challenge with the same strain (Figure S1A). We confirmed that *Shigella*-infected mice spontaneously clear the primary infection and are free of culturable *Shigella* by day 30 post-primary infection. Since primary *Shigella* infection of mice requires pre-treatment with streptomycin, we used streptomycin-treated but uninfected mice as a negative control. Unexpectedly, both previously-infected and streptomycin only-treated mice were resistant to colonization upon secondary challenge with virulent *Shigella* (Figure S1B). We hypothesized that streptomycin-induced alterations to the microbiome render mice resistant to a subsequent *Shigella* infection. To attempt to eliminate the putative protective species from the microbiome of our mice, we rederived our *Nlrc4*^−/−^ mice at Taconic at their stringent Opportunist Free (OF) standard. However, even OF-rederived mice were resistant to *Shigella* 30 days after streptomycin-only treatment, as compared to untreated controls (Figure S1C).

To circumvent the lingering effects of streptomycin on *Shigella* colonization, we decided to use a different antibiotic, ampicillin, to establish the secondary infection (scheme in Figure 1A). This approach was successful: streptomycin-treated mice that were gavaged with PBS were susceptible 30 days later to a *Shigella* infection established with ampicillin preclearance (one day prior to infection) (Figure 1B). The PBS-treated mice (hereafter referred to as “naïve”) lost up to 15% of their starting body weight upon *Shigella* challenge (Figure 1B). By contrast, streptomycin-treated mice that were given a primary *Shigella* infection (hereafter referred to as “immunized”) were resistant 30 days later to a secondary ampicillin-facilitated *Shigella* infection and maintained their body weight (Figure 1B). Immunized mice also had fewer *Shigella* colony forming units (CFUs) in their intestinal epithelial cells (IECs) relative to naïve controls (Figure 1C). Consistent with these data, immunized mice maintained their ceca lengths compared to PBS control mice (Figure 1D). Furthermore, confocal imaging demonstrated less expulsion of epithelial cells in the ceca of immunized mice as compared to naïve controls (Figure 1E). Analysis of colonic tissue and feces revealed that prior *Shigella* exposure protected against the inflammatory response upon secondary challenge. Specifically, immunized mice had reduced levels of IL-1β, CXCL1 and MPO relative to naïve controls (Figure 1F-H). The levels of bacteria in the intestinal lumen were similar between *Shigella*-infected naïve and immunized mice; in fact, the luminal colonization was slightly but significantly higher in immunized mice (Figure 1I). These data indicate that prior *Shigella* exposure can protect against epithelial cell infection and pathology following a subsequent challenge in our *Nlrc4*^−/−^ mouse model.

**Figure 1.**
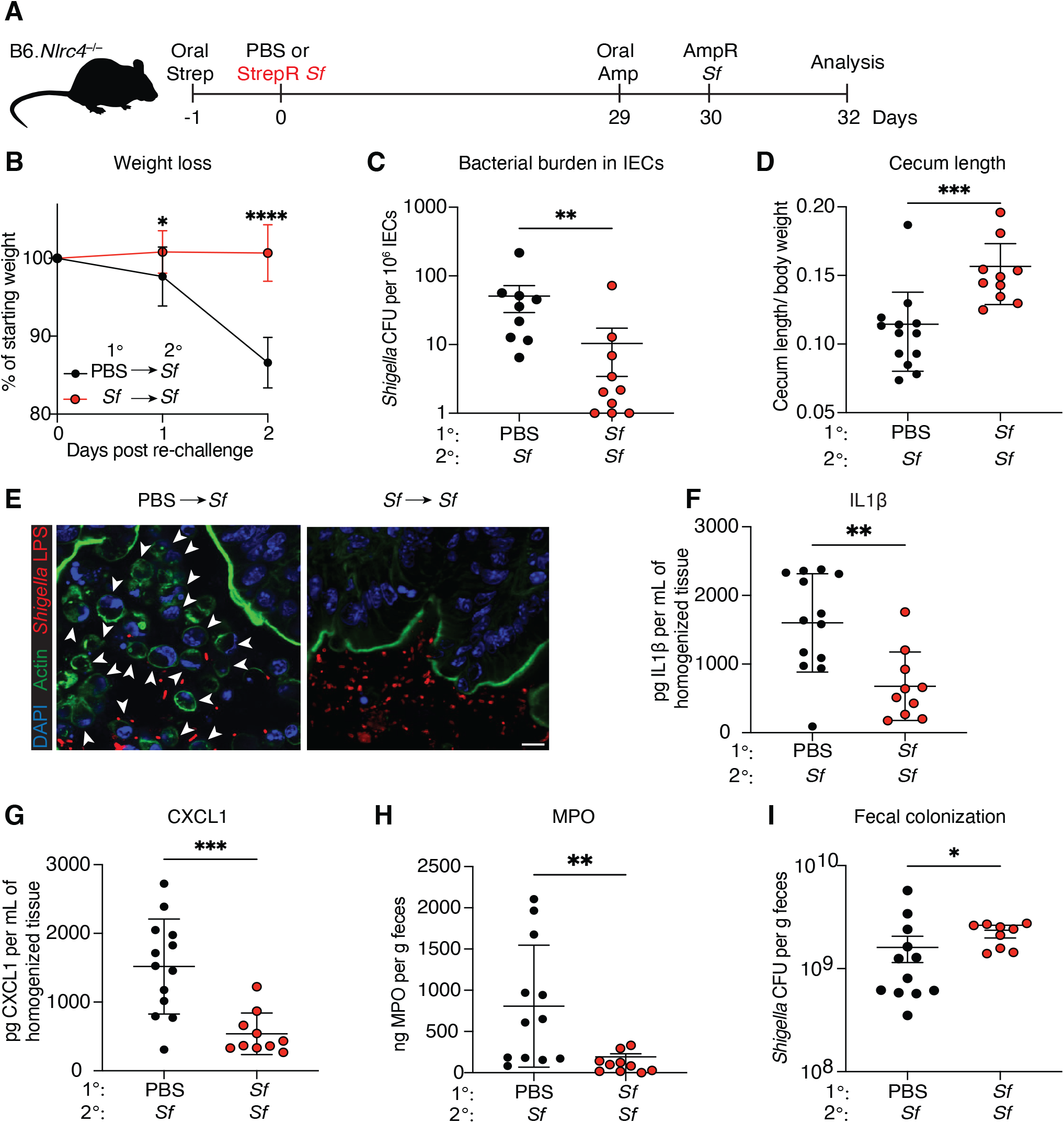
Primary *Shigella* infection protects against pathology upon re-challenge. (**A-I**) B6.*Nlrc4*^−/−^ mice were orally treated with 25mg streptomycin sulfate in PBS one day prior to oral challenge with 10^7^ colony forming units (CFU) of wild type (WT) *Shigella flexneri (Sf)* or PBS. Mice were left to recover for 28 days, followed by oral gavage with 20mg ampicillin sodium salt and re-challenge with 10^7^ CFU of WT *Sf* the next day. (**A**) Infection schematic. (**B**) Mouse weights from 0 to 2 days post re-challenge. Each symbol represents the mean (±SD) of mice of the indicated infection group. (**C**) *Shigella* CFUs per million intestinal epithelial cells. (**D**) Cecum length normalized to the mouse weight before re-infection. (**E**) Representative micrograph of cecal tissue from PBS control mice or mice previously infected with *Shigella* on re-infection with WT *Shigella*. White arrow heads represent expulsed IECs from the epithelial layer. (**F** and **G**) IL1 and CXCL1 levels measured by ELISA from homogenized colon and cecum tissue. (**H**) MPO levels measured by ELISA from homogenized feces. (**I**) *Shigella* CFU per gram of feces. Data are pooled from two independent experiments. Each symbol represents an individual mouse (C-I). Mean ±SD is shown in (B, C-H). Mean ±SEM is shown in (C and I). Statistical significance was calculated by two-way ANOVA with Šidák’s multiple comparison test (B), Mann-Whitney test (C and I) and Welch’s t test (D, F, G and H). *p<0.05, **p<0.01, ***p<0.001, ****p<0.0001, ns = not significant (p>0.05).

### Adaptive immunity is essential for protection from *Shigella* re-challenge

Next, we investigated the specific contribution of adaptive immune cells to protective immunity against secondary *Shigella* challenge. We crossed B6.*Nlrc4*^−/−^ mice with *Recombination activating gene 2* deficient mice (*Rag2*^−/−^) mice to generate *Nlrc4*^−/−^*Rag2*^−/−^ mice that lack mature B and T lymphocytes^42^. Co-housed *Nlrc4*^−/−^*Rag2*^−/−^ and *Nlrc4*^−/−^*Rag2*^+/+^ control mice were infected as in Figure 1A. *Nlrc4*^−/−^*Rag2*^−/−^ mice exhibited weight loss with kinetics comparable to *Nlrc4*^−/−^*Rag2*^+/+^ controls upon primary infection, with both groups reaching peak weight loss at day 3 post-infection and recovering by day 5 (Figure S2A). Both *Nlrc4*^−/−^*Rag2*^−/−^ and *Nlrc4*^−/−^*Rag2*^+/+^ mice cleared *Shigella* from their feces with similar kinetics (Figure S2B). These results indicate that adaptive immunity is not required for resolution of a primary *Shigella* infection. However, upon re-challenge, *Nlrc4*^−/−^*Rag2*^−/−^ mice displayed significant weight loss and had markedly higher *Shigella* bacterial burden in their IECs relative to *Nlrc4*^−/−^*Rag2*^+/+^ controls (Figure 2A, B). Consistent with these findings, *Nlrc4*^−/−^*Rag2*^−/−^ mice exhibited exacerbated pathology and inflammation, characterized by shortening of the ceca and elevated levels of IL1β, CXCL1 and MPO (Figure 2C-F). Collectively, these data reveal that adaptive immune cells are required for protection against bacterial colonization and pathology upon re-infection.

**Figure 2.**
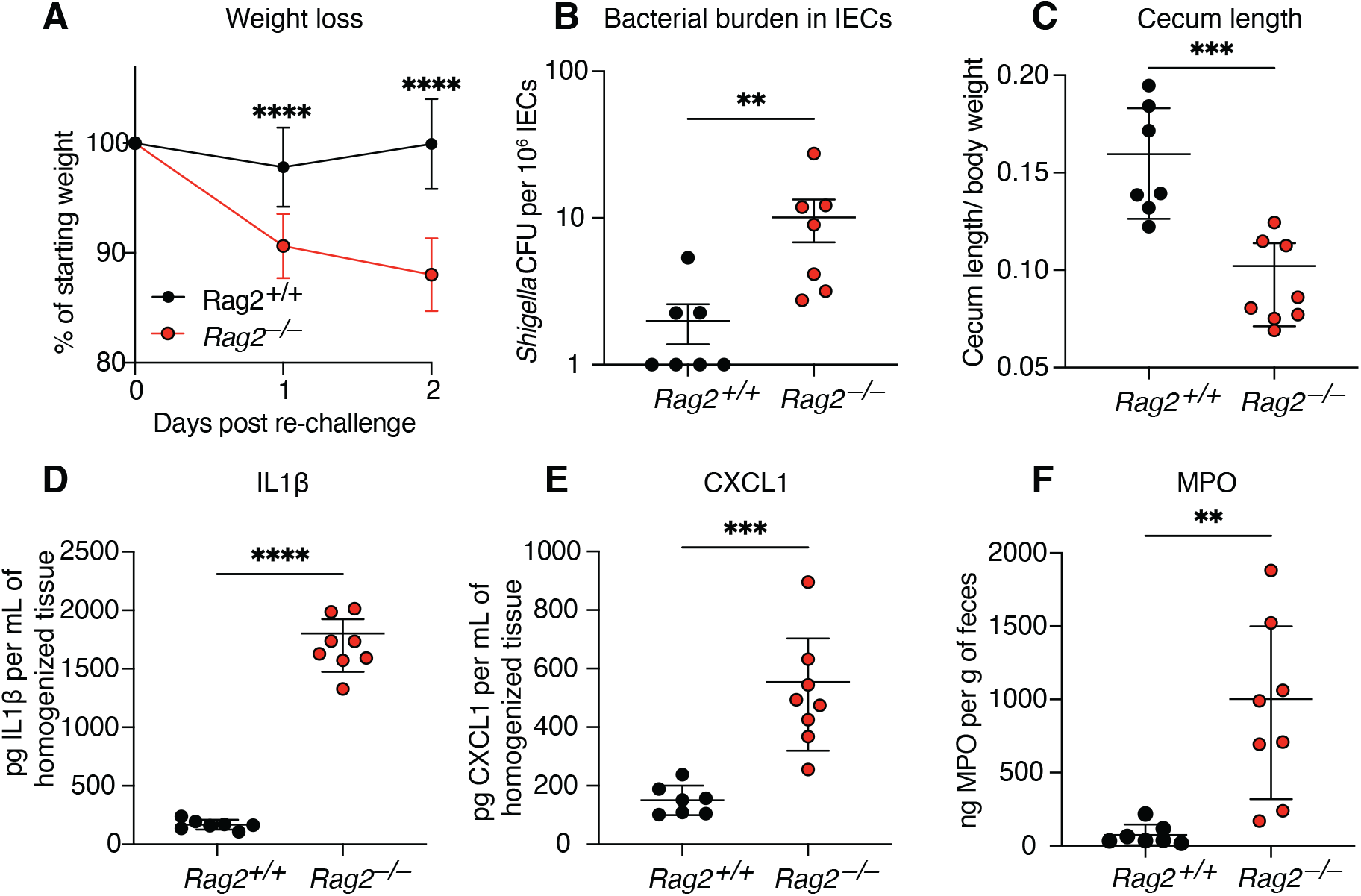
Adaptive immunity is required for protection against secondary challenge. (**A-F**) B6.*Nlrc4*^−/−^ or B6.*Nlrc4*^−/−^ *Rag2*^−/−^ mice were infected and re-challenged with WT *Shigella flexneri* as outlined in Figure 1. (**A**) Mouse weights from 0 to 2 days post re-challenge. Each symbol represents the mean (±SD) of mice of the indicated infection group. (**B**) *Shigella* CFUs per million intestinal epithelial cells on re-infection. (**C**) Cecum length normalized to the mouse weight before re-infection. (**D, E**) IL1b and CXCL1 levels measured by ELISA from homogenized colon and cecum tissue. (**F**) MPO levels measured by ELISA from homogenized feces. Data are representative of two independent experiments. (**B-F**) Each symbol represents one mouse. Mean ±SD is shown in (**A, C-F**). Mean ±SEM is shown in (B). Statistical significance was calculated by two-way ANOVA with Šidák’s multiple comparison test (A), Mann-Whitney test (B) and Welch’s t test (C-F). *p<0.05, **p<0.01, ***p<0.001, ****p<0.0001, ns = not significant (p>0.05).

### The *Shigella* virulence plasmid, but not cell-to-cell spread, is required to elicit protective immunity

Next, we sought to determine whether the *Shigella* virulence plasmid and/or cell-to-cell spread were required for the generation of protective immunity. To this end, we utilized two attenuated isogenic derivatives of *S. flexneri* 2457T: BS103, which lacks the virulence plasmid, and Δ*ics*A, which lacks the ability to hijack host actin to spread from cell to cell^15,43^. Both strains are attenuated in *Nlrc4*^−/−^ mice^33^. Mice were infected with either WT, BS103 or Δ*ics*A *S. flexneri*, and were then subsequently challenged with WT *S. flexneri* 2457T (as in Figure 1A). Mice previously infected with Δ*ics*A were protected from disease like WT-immunized mice, whereas mice previously infected with BS103 were susceptible to disease upon re-challenge with virulent *Shigella*. Mice immunized with Δ*icsA* were protected from weight loss, whereas the BS103 group exhibited significant weight loss, with kinetics mirroring those of naïve controls (Figure S3A). Moreover, significantly fewer bacterial CFUs were recovered from the IECs of Δ*icsA*-immunized mice as compared to mice previously infected with BS103 (Figure S3B). Mice previously infected with Δ*icsA* also maintained longer ceca, and displayed reduced tissue inflammation, as evidenced by significantly lower levels of IL-1β, CXCL1 and MPO, as compared to BS103-immunized or naïve controls (Figure S3C-F). Taken together, these data demonstrate that elicitation of protective immunity by *Shigella* requires the virulence plasmid but is independent of the ability of *Shigella* to spread cell-to-cell via actin-based motility.

### Oral *Shigella* infection induces robust CD4^+^ and CD8^+^ T cell responses

We next sought to characterize the T cell response against *Shigella* following oral infection. Because the specific *Shigella* antigens that activate endogenous T cell responses are unknown, we engineered *Shigella* to express the model antigens ovalbumin (Ova) and 2W to facilitate *in vivo* tracking of CD8^+^ and CD4^+^ T cell responses, respectively^44,45^. WT *Shigella flexneri* 2457T was transformed with a plasmid encoding FLAG-tagged 2W and Ova fused to the secretion signal of the OspC2 effector (Sf 2W-Ova), enabling antigen secretion via the T3SS (Figure S4A). *Shigella* transformed with an empty vector (Sf EV) served as a negative control.

To assess whether CD4^+^ T cell responses are generated upon primary *Shigella* infection, *Nlrc4*^−/−^ mice were streptomycin-treated and then infected orally with either Sf 2W-Ova or Sf EV. As a control, mice were also IP-injected with 2W peptide plus polyI:C as an adjuvant. At 7 days post-infection, splenocytes and mesenteric lymphocytes were isolated and analyzed for expansion of 2W-specific CD4^+^ T cells using 2W1S:I-A^b^ tetramers. We observed a significant increase in frequency and absolute number of 2W-specific CD4^+^ T cells in both the spleen and mLN of mice infected with Sf 2W-Ova compared to Sf EV controls (Figure 3A-C). Characterization of these 2W-specific CD4^+^ T cell subsets revealed expansion of Th1 (Tbet^+^), Th17 (Ror*γ*^+^) and Tfh (CXCR5^+^) helper subsets, but not FoxP3^+^ regulatory T cells (Figure 3D, Figure S4B).

**Figure 3.**
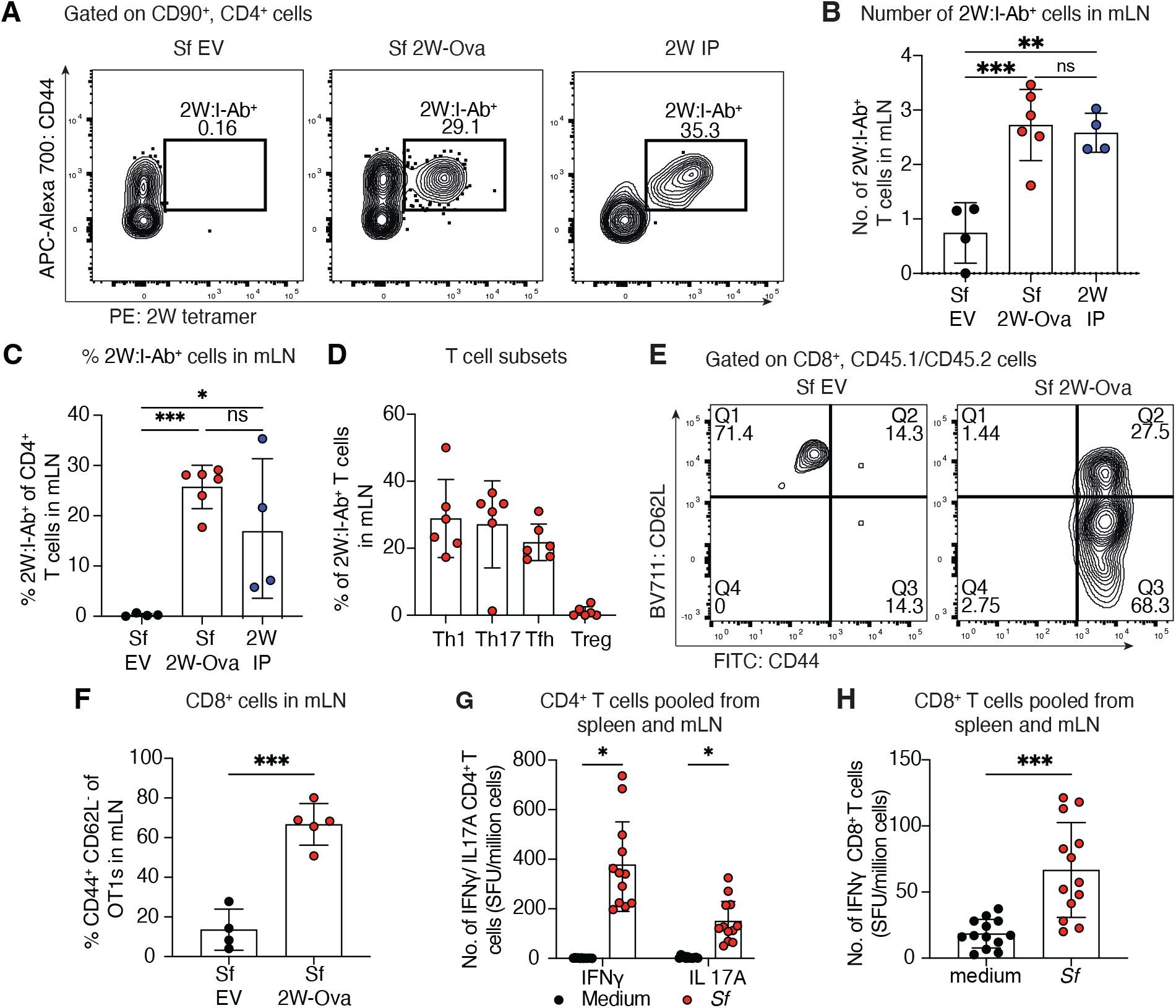
Primary Shigella infection induces CD4^+^ and CD8^+^ T cell expansion and activation. (**A-D**) Mice were infected with *Shigella* expressing 2W and Ova antigens (*Sf* 2W-Ova) or empty vector (*Sf* EV). Mice intraperitoneally injected with 2W peptide and polyIC (IP) served as a positive control. Splenocytes and mesenteric lymphocytes (mLN) were obtained and magnetically enriched for 2W^+^ CD4^+^ T cells using a tetramer and then analyzed by flow cytometry at 7 days post infection. (**A**) Representative flow plots indicate the frequency of 2W^+^ CD4^+^ T cells in mLN of mice 7d after receiving *Sf* EV, *Sf* 2W Ova or 2W IP. (**B**) Absolute number of 2W^+^ CD4^+^ T cells in mLN and (**C**) Percentage 2W^+^ of CD4^+^ T cells in mLN (**D**) Transcriptional factor expression profiling of 2W^+^ CD4^+^ T cells for Tbet^+^ (Th1), Ror*γ*^+^ (Th17), CXCR5^+^, (Tfh) and FoxP3^+^ (T regs). (**E-F**) OT1 cells (CD45.1/CD45.2) from a transgenic mouse were retro-orbitally transferred into B6.*Nlrc4*^−/−^ recipient mice (CD45.2). Mice were orally challenged with *Sf* 2W-Ova or *Sf* EV. (**E**) Representative flow plots for frequency of activated OT1 cells in recipient mice 7 days post. (**F**) Percentage of activated OT1s in mLN. (**G-H**) Mice were infected as in Figure 1 with WT *Shigella*. Splenocytes and mesenteric lymphocytes (mLN) were obtained and magnetically enriched for CD4^+^ T cells or CD8^+^ T cells. T cells were re-stimulated with irradiated splenocytes pulsed with heat-killed (HK) *Shigella*, and the number of IFN*γ* or IL-17 producing T cells enumerated by ELISpot assay as spot forming units (SFU). (**G**) Number of IFN*γ* and IL-17 producing CD4^+^ T cells upon restimulation with HK *Shigella*. (**H**) Number of IFN*γ* producing CD8^+^ T cells upon restimulation with irradiated *Shigella*. Mean ±SD is shown in (B, C, D, F, G and H). Statistical significance was calculated by ordinary one-way ANOVA (B and C), Welch’s t test (F and H) and multiple unpaired t tests (G). *p<0.05, **p<0.01, ***p<0.001, ****p<0.0001, ns = not significant (p>0.05)

To assess the CD8^+^ T cell response, Cell Trace Violet labeled Ova-specific CD8^+^ T cells (OT-I cells) were adoptively transferred into B6.*Nlrc4*^−/−^ mice one day prior to oral challenge with Sf 2W-Ova or Sf EV. Spleens and mLNs were analyzed at 7 days post infection by flow cytometry. Transferred OT-I cells proliferated in Sf 2W-Ova infected mice relative to Sf EV controls (Figure S4C-F), and upregulated CD44 and downregulated CD62L, confirming transition to the effector phenotype (Figure 3E and F).

To determine whether *Shigella* infection activates endogenous *Shigella*-specific T cells, we performed enzyme linked immunosorbent spot (ELISpot) assays on CD4^+^ or CD8^+^ T cells isolated from previously infected mice following *in vitro* restimulation with irradiated splenocytes and heat-killed *Shigella*. Consistent with the Sf 2W-Ova data, restimulation with heat-killed *Shigella* led to a significant increase in IFN*γ* and IL17A producing CD4^+^ T cells relative to medium controls (Figure 3G). Similarly, a higher frequency of CD8^+^ T cells from *Shigella* infected mice secreted IFN*γ* upon restimulation with irradiated *Shigella* (Figure 3H). Together, these results demonstrate that oral *Shigella* infection induces robust activation of both CD4^+^ and CD8^+^ T cells.

### CD4^+^ T cells, but not CD8^+^ T cells are required for adaptive immunity to *Shigella*

To dissect the contribution of CD4^+^ and CD8^+^ T cells during oral infection, we depleted either CD4^+^ or CD8^+^ T cells during primary *Shigella* infection to prevent the establishment of memory T cell responses (Figure 4A). T cell depletion was confirmed in peripheral blood on the final antibody administration via flow cytometry (Figure S5A, B). Upon secondary challenge, mice receiving anti-CD8 depleting-antibodies maintained body weight similar to isotype control IgG2b-treated mice. By contrast, anti-CD4-treated mice lost immune protection, and exhibited significant weight loss (Figure 4B). CD4^+^ but not CD8^+^ T cells were also required to prevent bacterial replication in IECs (Figure 4C). Furthermore, CD4^+^ T cell depletion resulted in exacerbated pathology and issue inflammation following *Shigella* re-challenge (Figure 4D-G). These data show that adaptive immunity elicited by *Shigella* depends on CD4^+^ T cells but does not require CD8^+^ T cells.

**Figure 4.**
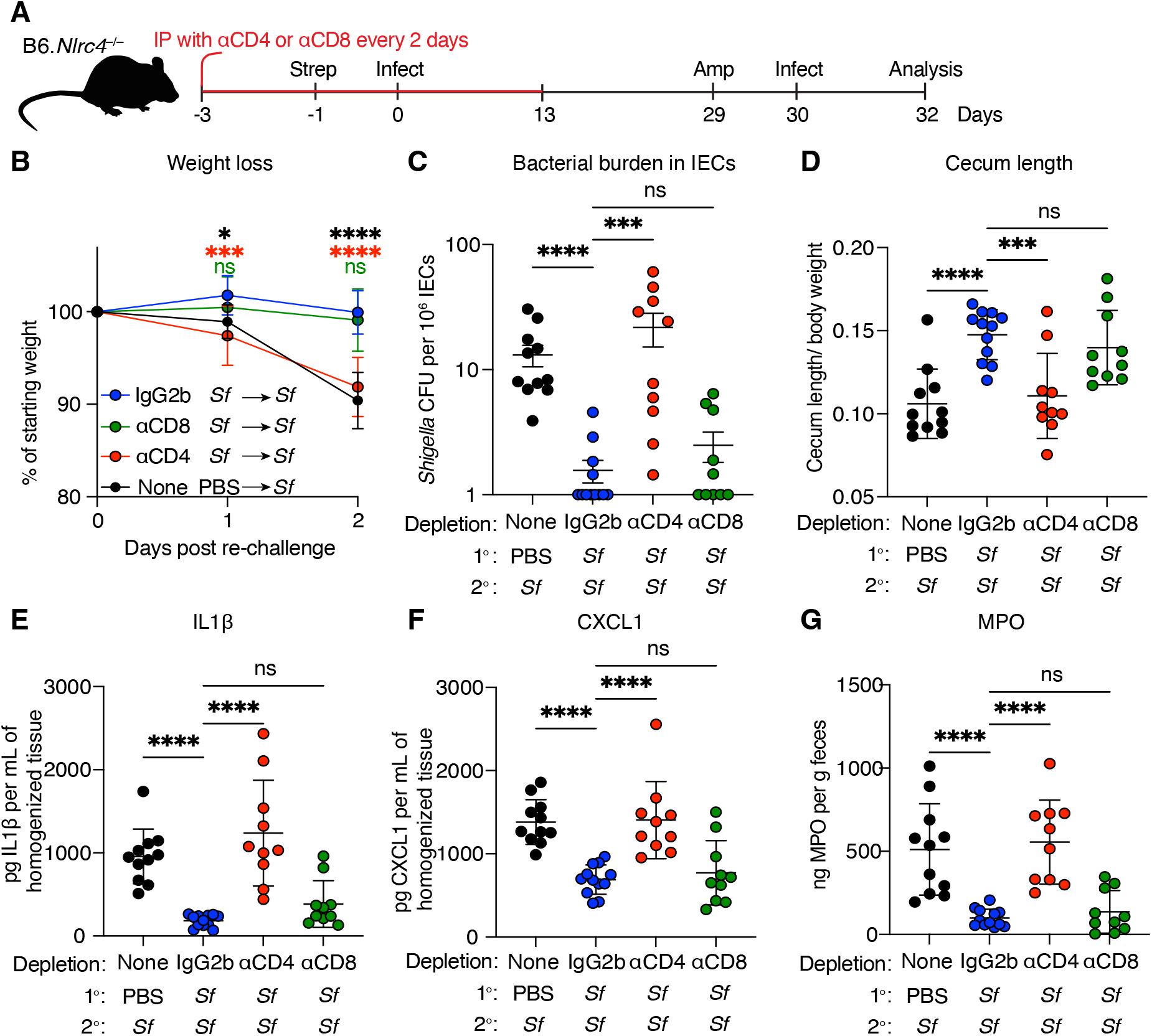
CD4^+^ T cells, but not CD8^+^ T cells are required for protection against secondary *Shigella* challenge. (**A-G**) Mice were IP injected with anti-CD4 (αCD4) or anti-CD8 (αCD8) depleting antibodies 3 days prior to primary infection with WT *Shigella*, and every 2 days post-infection for 2 weeks. Control mice received an IgG2b isotype control antibody. Mice were rechallenged with WT *Shigella* and sacrificed 2 days post re-infection. (**A**) Infection schematic. (**B**) Mouse weights from 0 to 2 days post re-challenge. Each symbol represents the mean of mice (±SD) of the indicated infection group. (**C**) *Shigella* CFUs per million intestinal epithelial cells. (**D**) Cecum length normalized to the mouse weight before re-infection. (**E, F**) IL1b and CXCL1 levels measured by ELISA from homogenized colon and cecum tissue. (**G**) MPO levels measured by ELISA from homogenized feces. Data are pooled from two independent experiments. Each symbol represents one mouse (**C-G**). Mean ± SD is shown in (B, D-G). Mean ± SEM is shown in (C). Statistical significance was calculated by two-way ANOVA with Tukey’s multiple comparison test (B), Kruskal-Wallis test (C) and ordinary one-way ANOVA (D-G). *p<0.05, **p<0.01, ***p<0.001, ****p<0.0001, ns = not significant (p>0.05).

### *Shigella* elicits an antibody response that is not essential for protection

To determine if primary *Shigella* infection induces an antibody response, naïve and immunized *Nlrc4*^−/−^ were re-challenged with mCherry*-*expressing *Shigella*^46^. IgA coating of fecal mCherry^+^ *Shigella* were quantified by microbiota flow cytometry^47^. *Shigella* in the feces of immunized (but not naïve) mice was coated with high levels of IgA (Figure 5A and B). We also observed increased levels of *Shigella* specific IgG in sera of immunized mice compared to naïve mice (Figure S6A and B). To determine if the anti-*Shigella* IgA response was T cell dependent, mice were treated with CD4^+^ depleting antibody or isotype control during the primary infection and subsequently re-challenged with mCherry^+^ *Shigella*. CD4^+^ T cell depletion reduced IgA coating of fecal *Shigella* to the level of naïve controls (Figure 5C, Figure S6C).

**Figure 5.**
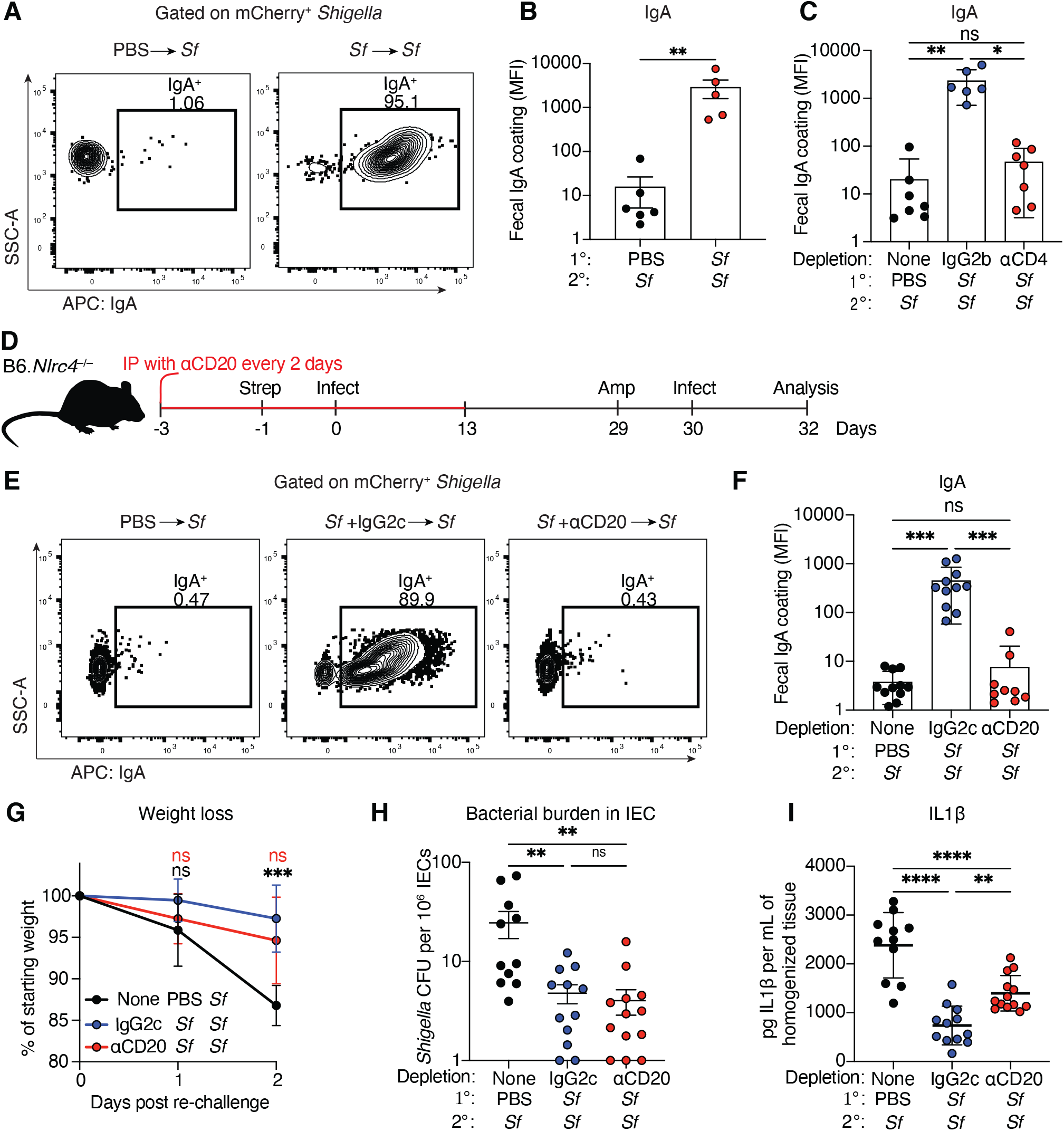
CD4-dependent IgA responses are elicited, but not necessary, for protective immunity against *Shigella*. (**A-C**) Mice were infected as outlined in Fig. 1 with WT *Shigella* or PBS. Mice were re-challenged with WT *Shigella* expressing mCherry. Feces were obtained and analyzed for IgA coating of *Shigella* by flow cytometry. (**A**) Representative flow plots for IgA coating of *Shigella* in feces. (**B-C**) Quantification of IgA MFI coating mCherry^+^ *Shigella* in undepleted (B) or CD4-depleted mice (C). (**D-J**) Mice were intraperitoneally administered with anti-CD20 depleting antibody 3 days prior to primary infection with WT *Shigella* and every 2 days post infection for 2 weeks. Control mice received an IgG2b isotype control antibody. Mice were re-challenged with WT *Shigella*. (**D**) Infection schematic. (**E**) Representative flow plots for IgA coating following B cell depletion. (**F**) Quantification of (E). (**G**) Mouse weights from 0 to 2 days post re-challenge. Each symbol represents the mean of mice of the indicated infection group (**H**) *Shigella* CFUs per million intestinal epithelial cells on re-infection. (**I**) IL1b levels measured by ELISA from homogenized colon and cecum tissue. Data are pooled from two independent experiments. Mean ±SD is shown in (B, C, F, G and I). Mean ±SEM is shown in (H). Statistical significance was calculated by Mann-Whitney test (B), Kruskal-Wallis test (C, F and H), ordinary one-way ANOVA (I) and two-way ANOVA with Tukey’s multiple comparison test (G). *p<0.05, **p<0.01, ***p<0.001, ****p<0.0001, ns = not significant (p>0.05).

We next sought to investigate whether B cells are required for immune protection against *Shigella*. B cells were depleted with anti-CD20 during the primary infection to prevent the development of *Shigella* specific B cell responses (Figure 5D). As expected, anti-CD20 antibody eliminated IgA coating of fecal *Shigella* during re-challenge (Figure 5E and F). However, immunized anti-CD20-treated mice were similar to immunized isotype-treated control mice, and exhibited less weight loss, lower bacterial burdens in IECs, and lower levels of pro-inflammatory IL-1β, compared to naïve controls (Figure 5G-I). Interestingly, immunized anti-CD20-depleted mice exhibited slightly higher levels of IL-1β than isotype control-treated mice, suggesting that B cells may provide modest protection against *Shigella*-induced inflammation. Taken together, these results demonstrate that oral *Shigella* infection induces a robust IgA response, but B cells are not essential for protection against subsequent infection.

### Cross-serotype immunity to *Shigella*

*Shigella flexneri* strains are subdivided into distinct serotypes based on structural variation in their lipopolysaccharide (LPS) O-antigens^48^. The above studies employ *Shigella flexneri* 2457T, a serotype 2a strain. To extend our observations to another serotype, we infected mice with *S. flexneri* M90T, a representative of the distinct 5a serotype. Interestingly, naïve mice infected with M90T exhibited less weight loss, reduced bacterial colonization, and diminished inflammation compared to naïve mice infected with 2457T (Figure S7A-F). There was decreased luminal colonization in mice infected with M90T compared 2457T (Figure S7G). Moreover, *Shigella* M90T isolated from the feces of infected mice exhibited a reduced frequency of Congo Red positivity, indicating loss of the virulence plasmid, which may explain the reduced virulence of this strain in mice (Figure S7H).

Despite its reduced virulence, we tested whether primary M90T infection nevertheless confers cross-protection against a subsequent 2457T challenge (Figure 6A). Mice immunized with M90T maintained their baseline weight following challenge with 2457T, similar to 2457T-immunized mice (Figure 6B). M90T and 2457T immunized mice exhibited similarly reduced *Shigella* CFUs in IECs compared to naïve controls (Figure 6C). Additionally, both M90T- and 2457T-immunized mice challenged with 2457T were protected from cecal shrinkage and had significantly lower levels of pro-inflammatory cytokines within the colonic tissues compared to PBS controls (Figure 6D-F). Thus, we observe cross-serotype protection in our *Nlrc4*^−/−^ mouse model of shigellosis.

**Figure 6.**
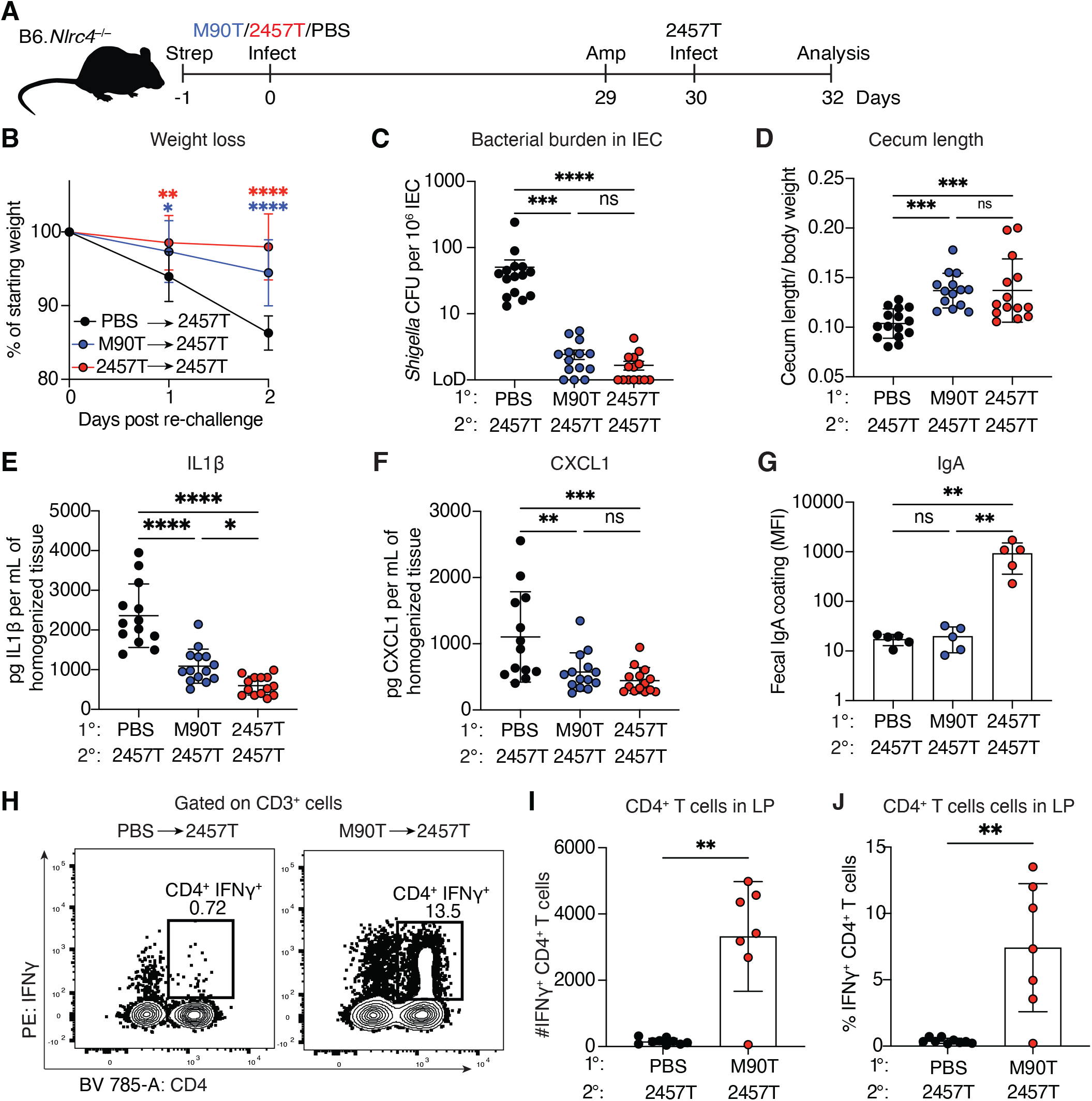
Cross-serotype protection is elicited by *Shigella flexneri* M90T. (**A-E**) Mice were infected with M90T (*S. flexneri* 5a) or 2457T (*S. flexneri* 2a) and subsequently re-challenged with 2457T. (**A**) Infection schematic (**B**) Mouse weights from 0 to 2 days post re-challenge. Each symbol represents the mean (±SD) of mice of the indicated infection group. (**C**) *Shigella* CFUs per million intestinal epithelial cells in re-infected mice. (**D**) Cecum length normalized to mouse weight before re-infection. (**E** and **F**) IL-1b and CXCL1 levels measured by ELISA from homogenized colon and cecum tissue. (**G**) *Shigella* specific IgA against 2457T. (**H-J)** Mice were infected with M90T (*S. flexneri* 5a) or 2457T (*S. flexneri* 2a) and subsequently challenged with 2457T. Mice were intravenously injected with brefeldin A 1 day post re-infection with 2457T and IFN*γ*-producing CD4^+^ T cells in the lamina propria were quantified. (**H**) Representative flow plots of IFN*γ*-producing CD4^+^ T cells in the lamina propria. (**I**) Number and (**J**) frequency of IFN*γ*-producing CD4^+^ T cells in the lamina propria (LP). Data are pooled from two independent experiments. Mean ±SD is shown in (B, D-G, I and J). Mean ±SEM is shown in (C). Statistical significance was calculated by two-way ANOVA with Tukey’s multiple comparison test (B), Kruskal-Wallis test (C and G), ordinary one-way ANOVA (D-F), and Welch’s t test (I and J). *p<0.05, **p<0.01, ***p<0.001, ****p<0.0001, ns = not significant (p>0.05).

To elucidate the mechanism of cross-serotype protection, we assessed IgA coating of *Shigella* isolated from feces of M90T-immunized mice. We did not observe significant IgA coating of *Shigella* 2457T in the feces of previously M90T-immunized mice, whereas 2457T was robustly IgA-coated in the feces of 2457T-immunized mice (Figure 6G). The lack of detectable cross-reactive IgA is consistent with LPS O-antigen being a major component of the anti-*Shigella* IgA response. During the innate immune response, IFN*γ* plays a critical role in limiting intracellular *Shigella* replication within IECs^41^. We hypothesized that lamina propria CD4^+^ T cells might mediate cross-protection via IFN*γ* secretion. Consequently, we first assessed whether CD4^+^ T cells produce IFN*γ* upon 2457T rechallenge in M90T-immunized mice. One day after re-challenge, we observed a robust population of IFN*γ*-producing CD4^+^ T cells in the lamina propria of M90T-immunized mice, which was greater than the IFN*γ* response seen in naïve controls (Figure 6H-J). There was also a significant population of IFN*γ*-producing CD8^+^ T cells in these mice (Figure S8A-C). In summary, *Shigella* can elicit cross-serotype protection that is associated with T cell IFN*γ* responses, but not mucosal IgA production.

### IFN*γ* is required for cross-serotype immunity against *Shigella*

We next sought to investigate whether homologous and/or heterologous (cross-serotype) immunity requires IFN*γ*. We first asked if IFN*γ* is required during a homologous 2457T re-challenge in which we observe robust IgA coating of the bacteria (Figure 5). Naïve or 2457T-immunized mice were (re-)challenged with 2457T while simultaneously treated with an anti-IFN*γ* neutralizing antibody (Figure 7A). As previously described, naïve mice required IFN*γ* to control a primary *Shigella* infection (Figure 7B-E). By contrast, IFN*γ* blockade did not exacerbate infection in 2457T-immunized mice, as IFN*γ* neutralized immunized mice maintained their initial body weight and exhibited low bacterial burdens and inflammation, similar to isotype control-treated mice (Figure 7B-E). Thus, IFN*γ* is not required for immunity during homologous infection, where there is a robust anti-*Shigella* IgA response. We then evaluated whether IFN*γ* is required in the context of heterologous (cross-serotype) immunity where there is no detectable anti-*Shigella* IgA response. M90T-immunized mice were re-challenged with 2457T in the presence of anti-IFN*γ* or isotype control antibodies. Interestingly, neutralizing IFN*γ* abolished protective cross-serotype immunity elicited by M90T. M90T-immunized IFN*γ*-neutralized mice exhibited weight loss, elevated bacterial burdens and inflammation similar to naïve IFN*γ*-neutralized mice (Figure 7F-I). In sum, these data indicate that IFN*γ* is required for the cross-serotype immunity to *Shigella* observed in our mouse model.

**Figure 7.**
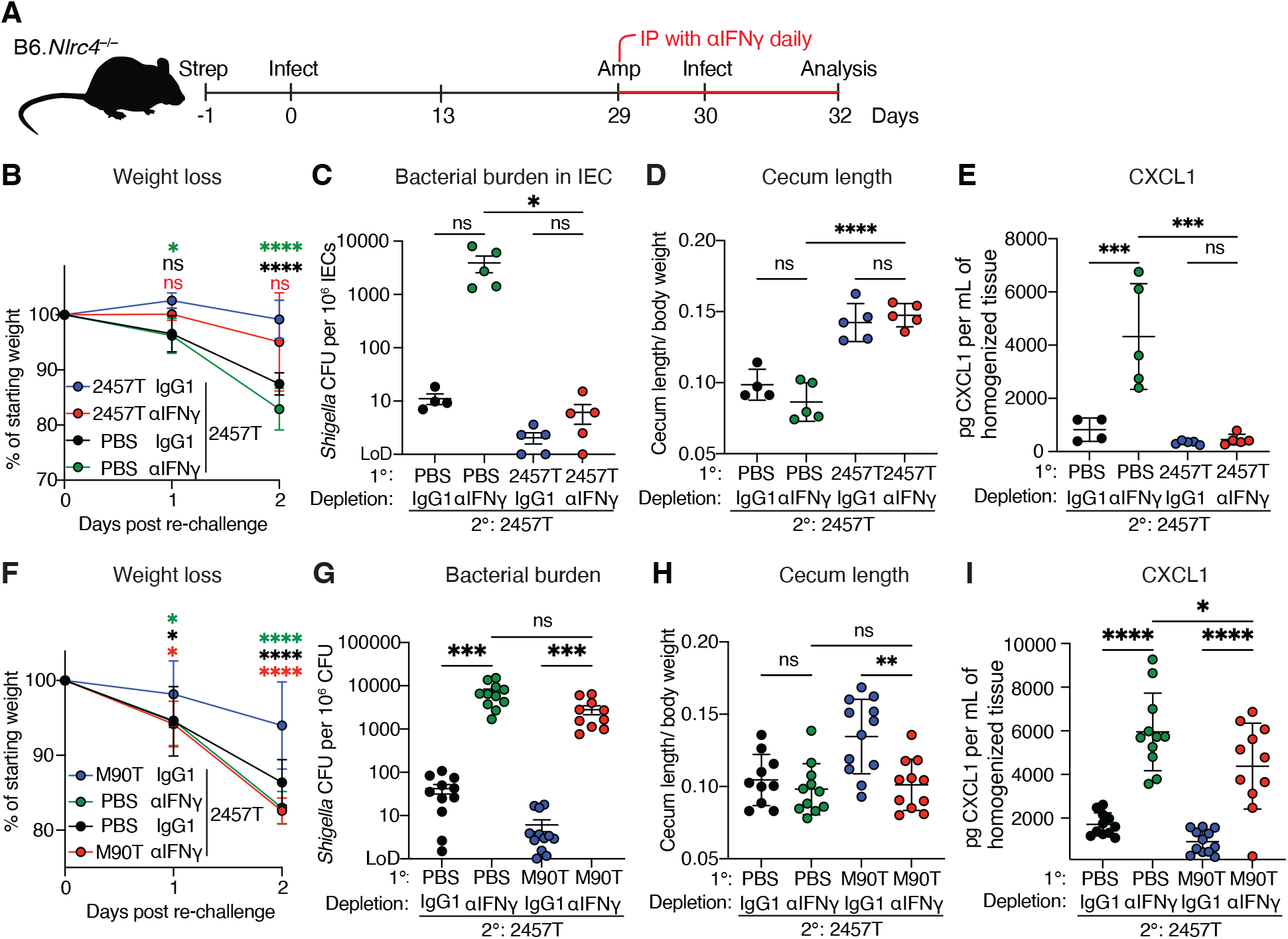
IFN*γ* is important for cross-serotype protection. (**A-I**) Streptomycin pre-treated mice were orally infected with 2457T (*S. flexneri* 2a) or M90T (*S. flexneri* 5a). After 28 days, mice were re-challenged with 2457T ± daily anti-IFN*γ* (αIFN*γ*) or isotype control (IgG1). (**B, F**) Mouse weights from 0 to 2 days post re-challenge. Each symbol represents the mean of mice of the indicated infection group. (**C, G**) *Shigella* CFUs per million intestinal epithelial cells on re-infection. (**D, H**) Cecum length normalized to the mouse weight before re-infection. (**E, I**) CXCL1 levels measured by ELISA from homogenized colon and cecum tissue. Data are pooled from two independent experiments. Mean ±SD is shown in (B, D-F, H and I). Mean ±SEM is shown in (C and G). Statistical significance was calculated by two-way ANOVA with Tukey’s multiple comparison test (B and F), Kruskal-Wallis test (C and G) and ordinary one-way ANOVA (D, E, H and I). *p<0.05, **p<0.01, ***p<0.001, ****p<0.0001, ns = not significant (p>0.05).

## Discussion

Decades of research have failed to arrive at a clear consensus as to which host responses correlate with protection against shigellosis. Even if such correlations were established, it would remain unclear whether any of these responses are causally protective. Causal studies of immunity require a physiological and experimentally tractable animal model, but the *Shigella* field has long lacked such a model. As a result, extensive ongoing efforts at developing an urgently needed *Shigella* vaccine are based on speculative assessments as to which responses the vaccine should elicit, and how they should be elicited. Such empirical approaches to vaccine development are common and can be successful, but it is preferable if they can be complemented with mechanistic studies that can explain why the vaccine works or fails, and how the vaccine can then be further optimized. Here we take advantage of our recent development of a physiological mouse model of shigellosis^33,34,41^ to establish a new and highly tractable experimental system that permits mechanistic studies of adaptive immunity to *Shigella*. Our mouse model employs the natural physiological route of oral infection and recapitulates all the key hallmarks of human shigellosis. In our study, we establish key experimental approaches and take foundational steps toward delineating the multilayered mechanisms of cellular and humoral adaptive immunity to *Shigella*.

We demonstrate that in our mouse model, as in humans, primary oral exposure confers protection against secondary challenge with the homologous strain (Figure 1B-H). As is also the case in humans^22,49^, we observe that an attenuated Δ*icsA* mutant strain can elicit protective immunity. Crucially, oral infection drives the robust activation and expansion of both CD4^+^ and CD8^+^ T cells (Figure 3A-H). This contrasts with previously used intranasal challenge mouse models, which fail to elicit CD8^+^ T cell responses^32^. Our results thus emphasize the necessity of physiological enteric priming to uncover the complete repertoire of adaptive immune responses to *Shigella*. We observe that activated CD4^+^ T cells undergo differentiation into Th1, Th17 and Tfh cell subsets (Figure 3D) consistent with a coordinated mucosal defense network of distinct cytokine and antibody responses.

We formally demonstrate that adaptive immunity is required for protection to a subsequent challenge (Figure 2A–F), and our cell depletion studies indicated that CD4^+^ T cells, but not CD8^+^ T cells, are required for immunity (Figure 4B-G). The observed dispensability of CD8^+^ T cells contrasts with the key role of cytotoxic CD8^+^ T lymphocytes in protection against other cytosolic bacterial pathogens such as *Listeria monocytogenes*^50–52^. One possible explanation for the apparent lack of a role for CD8^+^ T cells is that *Shigella* may subvert or evade CD8^+^ T cell-mediated immunity^32,53^. Any such impairment in the CD8^+^ T cell response would have to be selectively at the effector phase of the response since we see no defect in the activation and elicitation of *Shigella*-specific CD8^+^ T cells. It is also possible that CD8^+^ T cells do contribute to immunity to *Shigella*, but the contribution is not essential given additional redundant cellular and humoral responses.

In addition to our studies of cellular immunity, we also established methods to study the humoral response to *Shigella*. During secondary challenges of *Shigella*-immunized mice, we observed extensive IgA coating of enteric *Shigella in situ* (Figure 5A-C). *Shigella* infection of humans also results in a robust IgA response that has been suggested to correlate with protection^20,22,54^. Mucosal antibody could entrap or enchain bacteria^55^ or engage complement or neutrophil-mediated killing effector mechanisms^23^, impairing the ability of *Shigella* to invade intestinal tissue. However, B cell depletion failed to abrogate protection (Figure 5G-I), demonstrating that mucosal antibody responses are functionally redundant when an intact memory CD4^+^ T cell response is present. Similarly, heterologous challenge experiments revealed that mice immunized with strain M90T retained robust protection upon secondary challenge with strain 2457T, despite an absence of detectable cross-reactive mucosal IgA (Figure 6B-F). Instead, cross-serotype protection was accompanied by robust production of IFN*γ* by memory CD4^+^ T cells residing in the lamina propria (Figure 6H-J), and we found that IFN*γ* is required for cross-serotype immunity (Figure 7F-I). Interestingly, IFN*γ* was not required for immunity to a homologous challenge in which serotype-specific IgA responses are present (Figure 7B-E). Together, these findings suggest CD4^+^ T cells orchestrate protection via elicitation of a two-tiered mucosal cellular and humoral response. The first tier relies on B cell-derived mucosal IgA that limits bacterial invasion into the epithelium. Operating in parallel is a cross-serotype response driven by IFN*γ*, which directly restricts bacterial replication inside epithelial cells in the absence of serotype-specific antibodies.

Although we observe cross-serotype immunity in our model, there is little evidence for cross-serotype immunity in humans^19^, although definitive studies are lacking. Indeed, a major hurdle in the development of an effective *Shigella* vaccine is the extensive structural diversity of the bacterial lipopolysaccharide (LPS) O-antigen across circulating strains^56,57^. Leading candidate vaccines predominantly focus on O-antigen to elicit protective serum IgG^17,58^. However, even trivalent formulations encompassing the most prevalent serotypes cover only 64% of global circulating strains^59^. While strategies targeting highly conserved structural proteins, such as the invasion plasmid antigens (IpaB, IpaC and IpaD) that form the T3SS tip, offer a promising avenue to bypass serotype specificity, the cellular mechanisms governing cross-serotype immunity have remained ill-defined.

While we have established key methodological approaches for the study of both cellular and humoral responses to *Shigella*, and made several foundational observations, our study is just the first step in achieving a deeper mechanistic understanding of immunity to *Shigella*. Clearly there are many additional questions regarding the mucosal immune response to *Shigella* that remain to be addressed. One important contribution of our study is that we have established a physiologically relevant experimental system in which all these questions can now be answered. Together, our results add to established views on adaptive immunity to *Shigella* and suggest an expanded approach to benchmarking correlates of protection for vaccine candidates. While ongoing efforts focus on anti-LPS serum IgG antibodies as key indicators of protection^54,60–62^, our data reveal that CD4^+^ T cells provide a layered defense against *Shigella* re-challenge, via elicitation of mucosal IgA antibody and production of IFN*γ*. In general, elicitation of mucosal immune responses remains poorly understood^63,64^. Our results suggest *Shigella* now provides an excellent model with which to dissect the mechanisms of protective mucosal immunity.

## Materials and Methods

### Mice

*Nlrc4*^−/−^ mice were previously generated and described by Mitchell, Roncaioli et. al^33^. For this study, mice were re-derived, bred, and maintained at Taconic Biosciences (Germantown, New York, US) under opportunist free (OF) conditions that exclude Beta hemolytic Streptococcus (non-Group D), *Klebsiella oxytoca, Klebsiella pneumoniae, Pasteurella multocida*, Proteus spp., *Pseudomonas aeruginosa* and *Staphylococcus aureus*. The *Nlrc4*^−/−^*Rag2*^−/−^ mouse line was generated at Taconic by crossing *Nlrc4*^−/−^ animals to *Rag2*^−/−42^ background and was also maintained under OF conditions at Taconic. Mice were transferred from Taconic to the University of California, Berkeley, and housed under specific pathogen-free (SPF) conditions in accordance with protocols approved by the University of California, Berkeley Institutional Animal Care and Use Committee. Experiments utilized age-matched (8-12 weeks old) male and female mice. To control for microbiota variations, littermate controls were almost always used, or in rare instances, mice were co-housed for at least one week prior to primary infection.

### Bacterial strains

Experiments were conducted using a streptomycin-resistant derivative of *Shigella flexneri* serovar 2a (strain 2457T) and serovar 5a (strain M90T)^65,66^. Isogenic derivatives of 2457T included BS103, a virulence plasmid-cured strain, and a Δ*icsA* mutant strain. Ampicillin-resistant (AmpR) *Shigella* was generated by transforming strain 2457T with BASIC_5_Amp, a plasmid that confers ampicillin resistance (Addgene, Cat#68139). Fluorescent *Shigella* was generated by transforming strain 2457T with a plasmid expressing mCherry^46^. To construct a *Shigella* strain expressing the 2W (EAW-GALANWAVDSA) and Ova (SIINFEKL) epitopes, a DNA sequence was synthesized (IDT) that encodes the IpaH7.8 promoter upstream of an *E. coli* codon-optimized fusion protein consisting of the type III secretion signal sequence of OspC2, a flexible Gly-Ser linker, the 2W epitope, a fragment of chicken ovalbumin containing both the OT-I and OT-II epitopes, and a C terminal 3xFLAG tag, all flanked by attB1 sites. This sequence was inserted by Gateway cloning (Invitrogen) into the pDONR221 entry plasmid and then subsequently moved into pCMD^67^ (conferring spectinomycin resistance) containing attP sites. The resulting expression plasmid was isolated, sequence-verified, and subsequently transformed into 2457T.

### *Shigella* culture

*Shigella* was cultured at 37°C on Tryptic Soy Broth (TSB, BD Bacto, Cat# DF0370-07-5) agar plates supplemented with 0.01% Congo Red (CR, Sigma-Aldrich, Cat#C6767) and 100 μg/mL streptomycin sulfate. A single CR-positive colony from a fresh streak was inoculated into 5 mL of TSB containing 100 μg/mL streptomycin sulfate (and 100 μg/mL ampicillin sodium sulfate for AmpR *Shigella*) and grown overnight at 37°C with shaking. The overnight culture was diluted 1:100 into 5 mL of fresh TSB containing 100 μg/mL streptomycin sulfate and incubated at 37°C with shaking for approximately 3 hours until reaching an OD^600^ of 1.4. Bacteria were harvested by centrifugation at 5,000 ×*g* for 5 minutes, washed twice with PBS, and resuspended in PBS to a final concentration of 1×10^8^ CFU/mL.

### *In vivo* infections, depletions and Brefeldin A injections

One day prior to infection, mice were fasted for 4–6 hours and administered 25 mg of streptomycin sulfate or 20 mg of ampicillin sodium sulfate (in 100 μL sterile PBS) via oral gavage. The following day, mice were fasted again for 4–6 hours and orally gavaged with 100 μL of 1×10^8^ CFU/mL *Shigella* in PBS. Body weights were recorded daily, and any mice exhibiting greater than 20% weight loss relative to baseline were humanely euthanized in accordance with institutional guidelines.

For *in vivo* IFN*γ* neutralization, mice were intraperitoneally (i.p) injected daily during re-challenge with 500 μg (in 100 μL) of anti IFN*γ* antibody (clone XMG1.2 Bio X cell, Cat# BE055) or *In Vivo* MAb rat IgG1 isotype control, anti-horseradish peroxidase (clone HRPN, Bio X Cell, Cat# BE0088). For *in vivo* cell depletions, mice received an initial i.p. injection of 500 μg (in 100 μL) of anti-mouse CD20 (clone MB20-11, BioXCell, Cat#BE0356), anti-mouse CD4 (clone GK 1.5, Leinco technologies, Cat# C1333), or anti-mouse CD8 (clone YTS-169, Leinco technologies, Cat# C2442) antibodies 3 days prior to primary infection, followed by maintenance doses of 200 μg (in 100 μL) every 2 days for a total of 2 weeks. Control mice were treated on the same schedule with IgG2c isotype control, anti-dengue virus (clone DV5-1, BioXCell, Cat# BE0366; control for anti-CD20) or Rat IgG2b isotype control (clone 1-2, Leinco technologies, Cat# I-1034; control for anti-CD4 and anti-CD8).

To measure intracellular IFN*γ in vivo*, mice were injected intravenously via the lateral tail vein with 250 μg of Brefeldin A (Sigma, Cat# B6542) diluted in 100 μL of sterile PBS following brief warming under a heat lamp. At 6 h post injection, animals were euthanized and tissues were harvested as indicated below.

### Fecal CFUs and MPO ELISA

Fecal samples were collected on 1- and 2-days infection in pre-weighed 2 mL cryotubes and weighed to determine net stool mass. Samples were resuspended in 500 μL of PBS containing 2% FBS and protease inhibitors, then homogenized. For bacterial enumeration, homogenates were serially diluted in PBS and plated on TSB agar supplemented with 0.01% Congo Red and 100 μg/mL streptomycin sulfate. For MPO quantification, homogenates were centrifuged at 3,000 ×*g* for 3 minutes, and the resulting supernatants were analyzed using a mouse MPO ELISA kit (R&D Systems, Cat# DY3667) according to the manufacturer’s instructions.

### Tissue harvesting and IEC processing

Ceca and colons were harvested 2 days post-rechallenge as previously described^33^. Briefly, cecal and colon lengths were recorded before tissues were opened longitudinally to remove luminal contents. Tissues were washed in cold PBS and stored on ice for 1–2 hours in 5 mL of RPMI 1640 (Thermo Fisher Scientific, Cat# 21870) supplemented with 5% FBS, 2 mM GlutaMAX (Thermo Fisher Scientific, Cat# 35050061), 25 mM HEPES (Thermo Fisher Scientific, Cat# 15630080), and 400 μg/mL gentamicin (Thermo Fisher Scientific, Cat# 15710064). Samples were vortexed to remove residual fecal material and washed six times in cold PBS. Tissues were then cut into ~1 cm pieces and incubated in 12 mL stripping buffer (HBSS, 25 mM HEPES, 2 mM GlutaMAX, 50 μg/mL gentamicin, 2 mM DTT, and 5 mM EDTA) at 37 °C for 30 minutes with gentle agitation. The supernatant containing the IEC fraction was passed through a 100 μm cell strainer. Remaining tissue pieces were shaken vigorously in 10 mL of PBS, strained through the same filter to combine IEC fractions, and retained in 1mL PBS containing 2% FBS and protease inhibitors (Thermo Scientific, Cat# A32965) for tissue ELISAs. The isolated IEC fraction was incubated with 50 μg/mL gentamicin on ice for 20 minutes, pelleted by centrifugation (300 ×*g*, 7 min, 4 °C), and washed twice with ice-cold PBS. An aliquot (500 μL) was removed for cell quantification, and the remaining cells were lysed in 1 mL of 1% Triton X-100. Lysates (100 μL) were plated on TSB agar supplemented with 0.01% Congo Red and 100 μg/mL streptomycin sulfate to quantify intracellular bacteria.

### Lamina propria preparation

Colons and ceca were harvested as previously described, minced into small fragments and incubated in 12 mL stripping buffer (HBSS, 25 mM HEPES, 2 mM GlutaMAX, 50 μg/ mL gentamicin, 2 mM DTT, and 5 mM EDTA) for 30 minutes at 37 °C with continuous magnetic stirring (~400 rpm). The supernatant containing the IEC fraction was passed through a 100 μm cell strainer. Remaining tissue fragments were shaken vigorously in 10 mL of PBS, strained through the same filter to combine IEC fractions, and returned to the flask. Retained tissues were finely minced with scissors in microcentrifuge tubes and returned to 25 mL Erlenmeyer flasks containing 10 mL of lamina propria (LP) digestion buffer (HBSS supplemented with 1 mg/mL Collagenase VIII and 50 μg/mL DNase I). Samples were incubated for 25 min at 37 °C under 5% CO_2_ with continuous magnetic stirring (~400 rpm). Digestion was stopped on ice by adding 5 mL of cold RPMI-1640 supplemented with 10% FBS. Remaining fragments were mechanically dissociated with a syringe plunger, and flasks were rinsed with 10 mL cold RPMI-1640 with 10% FBS through the same strainer. Cells were centrifuged at 500 ×*g* for 5 min at 4°C, resuspended in 1 mL PBS containing 10% FBS, and layered onto a discontinuous Percoll density gradient (6 mL 37.5% Percoll in HBSS layered over 4 mL 70% Percoll in PBS). Gradients were centrifuged at 500 ×*g* for 8 min at room temperature without brakes. The upper layer was aspirated, and the enriched LP interphase collected, washed, and resuspended in HBSS with 5% FBS for downstream immunophenotyping. LP lymphocytes were stained with extracellular and intracellular antibodies (supplementary table 1) as previously described.

### Tissue ELISAs

Tissue reserved during IEC isolation was homogenized and centrifuged at 3,000 ×*g* for 5 minutes to collect supernatants. Protein concentrations of IL-1β and CXCL1 were quantified using ELISA kits (R&D Systems, Mouse CXCL1/KC DuoSet ELISA, Cat# DY453 and Mouse IL-1 beta/IL-1F2 DuoSet ELISA, Cat# DY401) according to the manufacturer’s instructions.

### Enzyme linked immunosorbent spot (ELISpot) assays for IFN*γ* and IL17A

ELISpot assays were performed as previously described^68^. Briefly, spleens and/or mesenteric lymph nodes (mLNs) were harvested from mice 7 days post-*Shigella*, infection. Tissues were mechanically dissociated between sterile frosted glass slides, filtered through a 100 μm nylon mesh, and centrifuged at 300 ×*g* for 7 min at 4°C. CD4^+^ or CD8^+^ T cells were purified via negative selection using magnetic enrichment T cell isolation kits (CD4^+^; Miltenyi, Cat# 130-104-454 or CD8^+^; Miltenyi, Cat#130-104-075) according to manufacturer’s protocol. ELISpot plates (Millipore, Cat#MAIPS4510) were coated overnight at 4°C with 100 μL/well of 15ug/mL anti-mouse IFN*γ* (clone AN-18, eBioscience, Cat#14-7313-85) or anti-mouse IL17A (BioLegend, Cat# 506902) capture antibody in sterile PBS. Plates were washed and blocked with complete RPMI supplemented with 10% FBS. Enriched CD4^+^ T or CD8^+^ T cells (2.5×10^5^ cells/well) were co-cultured with irradiated splenocytes (2.5×10^5^ cells/well) that had been pre-pulsed with heat killed or irradiated *Shigella*, or Concanavalin A (Sigma, Cat# C2010) as a positive control. Plates were incubated at 37°C with 5% CO_2_ for a minimum of 18 hours. Plates were washed with PBS and incubated with biotinylated anti-mouse IFN*γ* (clone R4-6A2, 1 μg/mL; eBioscience, Cat# 13-7312-85) or anti-mouse IL17A (1 μg/mL; BioLegend, Cat# 507002) detection antibody for 1.5 h at room temperature. Following additional PBS washes, streptavidin-alkaline phosphatase (Miltenyi Biotech, Cat#434322) diluted 1:1000 was added for 1 h in the dark. Spots were developed by adding one step NBT/BCIP substrate solution (Thermo Fisher Scientific; Cat#34042) for 5–15 min in the dark until spots were visible. The reaction was stopped by washing with deionized water and air-dried overnight. Spot forming units (SFU) were quantified using the immunospot reader.

### Microbial flow cytometry for IgA coated *Shigella*

Fecal pellets (1-2 per mouse) were collected from animals infected with mCherry-expressing *Shigella* and homogenized in 1mL PBS using a powerLyzer 24 homogenizer. Large particulate debris was removed by centrifugation at 250×*g* for 4 min. The supernatant was passed through a mesh-cap flow cytometry tube, diluted with 1mL of PBS, and centrifuged at 4,000×*g* for 5 min to pellet bacterial cells. The resulting bacterial pellet was resuspended in 1mL PBS, and the OD_600_ was measured to adjust the suspension to a final concentration of 5×10^7^ cells/mL in staining buffer (filtered PBS containing 2% BSA and 0.05% NaN_3_). A 25μL aliquot of the normalized bacterial suspension was incubated with 25μL of biotinylated anti-mouse IgA antibody (1:500; Southern biotech, Cat#1030-08) in a V-bottomed plate for 1h at 4°C. Plates were washed with 100μl staining buffer, centrifuged at 4,000×*g* for 5 min, and incubated with 50μl streptavidin-APC (1:1000 in PBS; Biolegend) in staining buffer for 30 minutes at 4°C in the dark. Following a final wash, bacteria were resuspended in 200μl of SYTOX BC nucleic acid stain at 1:1000 (ThermoFisher, Cat# S34855). Endogenous IgA coating of mCherry-expressing *Shigella* was subsequently quantified by flow cytometry.

### Cell preparation and adoptive transfer

Secondary lymphoid tissues (spleen and axial, brachial, inguinal, and mesenteric lymph nodes) were harvested from OT1 *Rag2*^−/−^ transgenic mice under sterile conditions and collected in complete RPMI on ice. Tissues were mechanically dissociated between sterile frosted glass slides, filtered through a 100 μm nylon mesh, and centrifuged at 300 x g for 7 min at 4°C. Erythrocytes were lysed by resuspending the cell pellet in 1 mL ACK Lysing Buffer (Gibco, Cat#A10492-01) for 5 min at room temperature, followed by quenching with 13 mL PBS and filtration through a 100 μm strainer. Splenocytes and lymphocytes were pelleted (300 x g for 7 at 4°C) and resuspended in PBS for cell counting. For fluorescent proliferation tracing, cells were adjusted to 1 × 10^6^ cells/mL in PBS and labeled with Cell Trace Violet (Thermo Fisher, Cat#C34557) according to the manufacturer’s instructions, keeping samples protected from light. Labeled cells were recounted and resuspended in PBS to a volume corresponding to the target cell dose in 100μL per recipient mouse. Recipient mice were briefly anesthetized with isoflurane and injected retro-orbitally with 100μL of the prepared cell suspension.

### Tetramer enrichment and flow cytometry

Tetramer staining was carried out as previously described^69^. Briefly, single-cell suspensions from harvested organs were washed in cold FACs buffer (PBS containing 2% FBS and 0.1% NaN_3_) by centrifugation at 300×*g* for 7 minutes at 4°C, and cell pellets were resuspended in 200 μL FACs buffer. Cells were stained with 2 μL of 1 μM 2W:I-A^b^/streptavidin-PE tetramer (NIH Tetramer Core), 4 μL anti-CXCR5 (1:50) and 2 μL Fc block (1:100) for 1 hour at room temperature in the dark. Cells were washed and resuspended in 500μL FACs buffer. Tetramer-bound cells were enriched using EasySep PE Positive Selection Kit II (Stem Cell technologies) by sequential incubations with 6.25 μL of EasySep PE Selection Cocktail (15 min) and 25 μL magnetic particles (10 min) at room temperature in the dark. Samples were brought to 2.5 mL with FACs buffer, filtered through nylon mesh, and loaded onto EasySep magnets for 5 min. Unbound fractions were discarded, and bound fractions were washed on the magnet three additional times before pelleting (600x g, 5 min at 4°C). Enriched cell pellets were surface stained with antibody cocktails (supplementary table 1) for 25 min at 4°C in the dark and fixed in 400 μL eBio-science Foxp3/Transcription Factor Fixation/Permeabilization solution (Thermo Fisher) for 45 min at room temperature in the dark. Cells were washed twice in 1× permeabilization buffer (1026×*g*, 5 min at 4 °C) and incubated overnight at 4°C in the dark with intracellular transcription factor antibodies (supplementary table 1) diluted 1:100 in permeabilization buffer. Finally, cells were washed with 3mL of FACs buffer (1000×*g*, 5 min at 4°C) and resuspended in 100 μL of FACS buffer for flow cytometric analysis.

### Microscopy

Cecal tips with intact luminal contents were fixed in Cytofix/Cytoperm (BD biosciences #554722) diluted 1:2 in PBS for 24h at 4°C. Tissues were washed with PBS and cryoprotected in 20% sucrose/PBS for 12h at 4°C. Cecal samples were embedded in OCT compound (Tissue-Tek) and stored at −80°C. Cryosections (10μm) were prepared on a Leica CM3050S cryostat and mounted onto Superfrost Plus slides (Fisher Scientific). Sections were rehydrated with PBS, permeabilized with 0.5% Triton X-100 in PBS, and blocked with 10% normal goat serum (Vector Laboratories) prior to staining with DAPI (Sigma) and fluorophore-coupled antibodies (supplementary table 1). Sections were mounted using Vectashield HardSet (Vector Laboratories) imaged on a Zeiss LSM710 confocal microscope. Image processing and analysis were performed using FIJI software.

### Statistical Analysis

Statistical tests to determine significance were performed using GraphPad prism software and are indicated in the figure legends.

## Acknowledgments

We thank Rebecca Brewer, Greg Barton, Jenna Vickery, and members of the Vance and Barton for their technical support and insightful discussions, and Gabriel Mitchell for comments on the manuscript. We are also grateful to UC Berkeley’s Flow Cytometry Facility (Cancer Research Laboratory), Biological Imaging Facility, and Office of Laboratory Animal Care for technical and facilities support.

## Author contributions

Conceptualization: J.P.B., S.A.F., C.F.L., and R.E.V.; investigation: J.P.B., E.L., S.Y., R.C., C.N., K.E., S.A.F. and D.I.K.; data analysis: J.P.B; methodology: J.P.B., S.A.F., D.I.K. Funding acquisition: R.E.V.; writing: J.P.B and R.E.V.

---

**Supplementary Figure 1.**
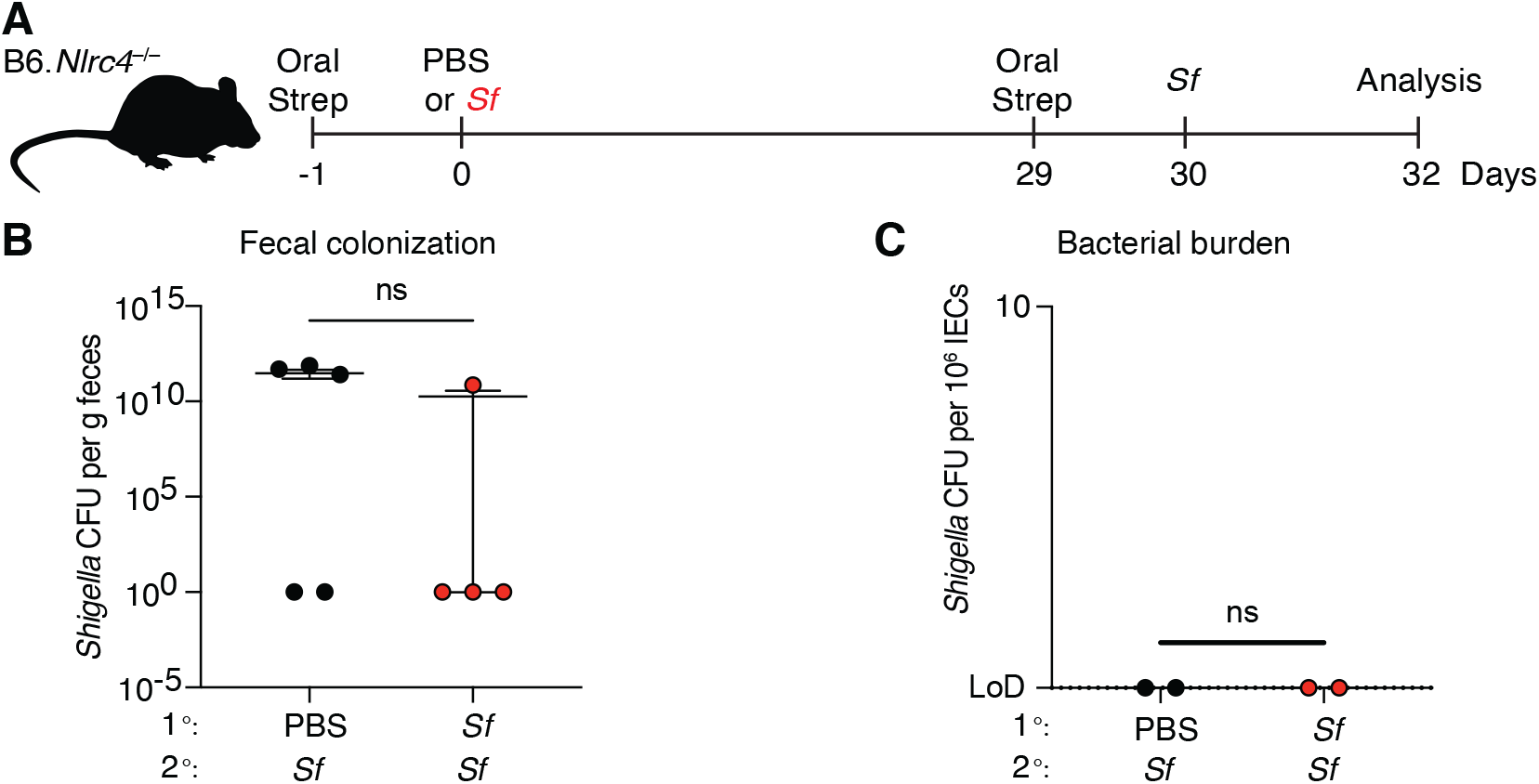
Primary streptomycin treatment renders mice resistant to secondary streptomycin treatment. (**A-B)** Mice were orally treated with streptomycin sulphate in PBS one day prior to oral challenge with 10^7^ CFU colony forming units (CFU) of wild type (WT) *Shigella flexneri (Sf)* or PBS. Mice were left to recover for 28 days, followed by oral gavage with streptomycin sodium salt and re-challenge with 10^7^ WT *Sf* the next day. (**A**) Infection schematic. (**B**) *Shigella* CFU per gram of feces. (**C**) Mice re-derived at Taconic were infected as described above and analyzed for bacterial CFU in IECs. (**C**) *Shigella* CFUs per million intestinal epithelial cells. Mean ±SEM is shown in (B-C). Mann-Whitney test (B-C). *p<0.05, **p<0.01, ***p<0.001, ****p<0.0001, ns = not significant (p>0.05).

**Supplementary Figure 2.**
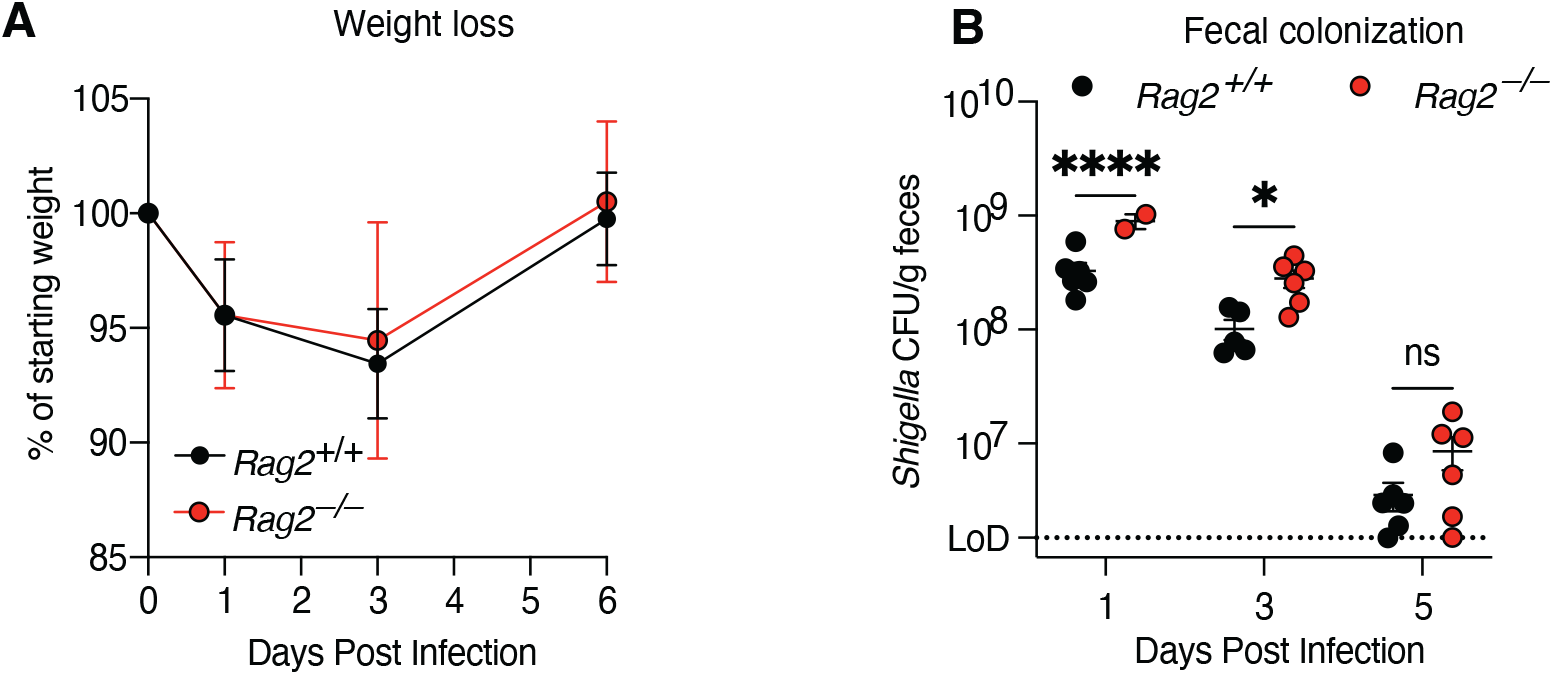
Adaptive immunity is not required for protection during primary infection. Mice were infected with *Shigella* and their weight loss and *Shigella* fecal CFUs monitored over time. (**A**) Mouse weights from 0 to 2 days post primary infection. Each symbol represents the mean (±SD) of mice of the indicated infection group. (**B**) *Shigella* CFU per gram of feces.

**Supplementary Figure 3.**
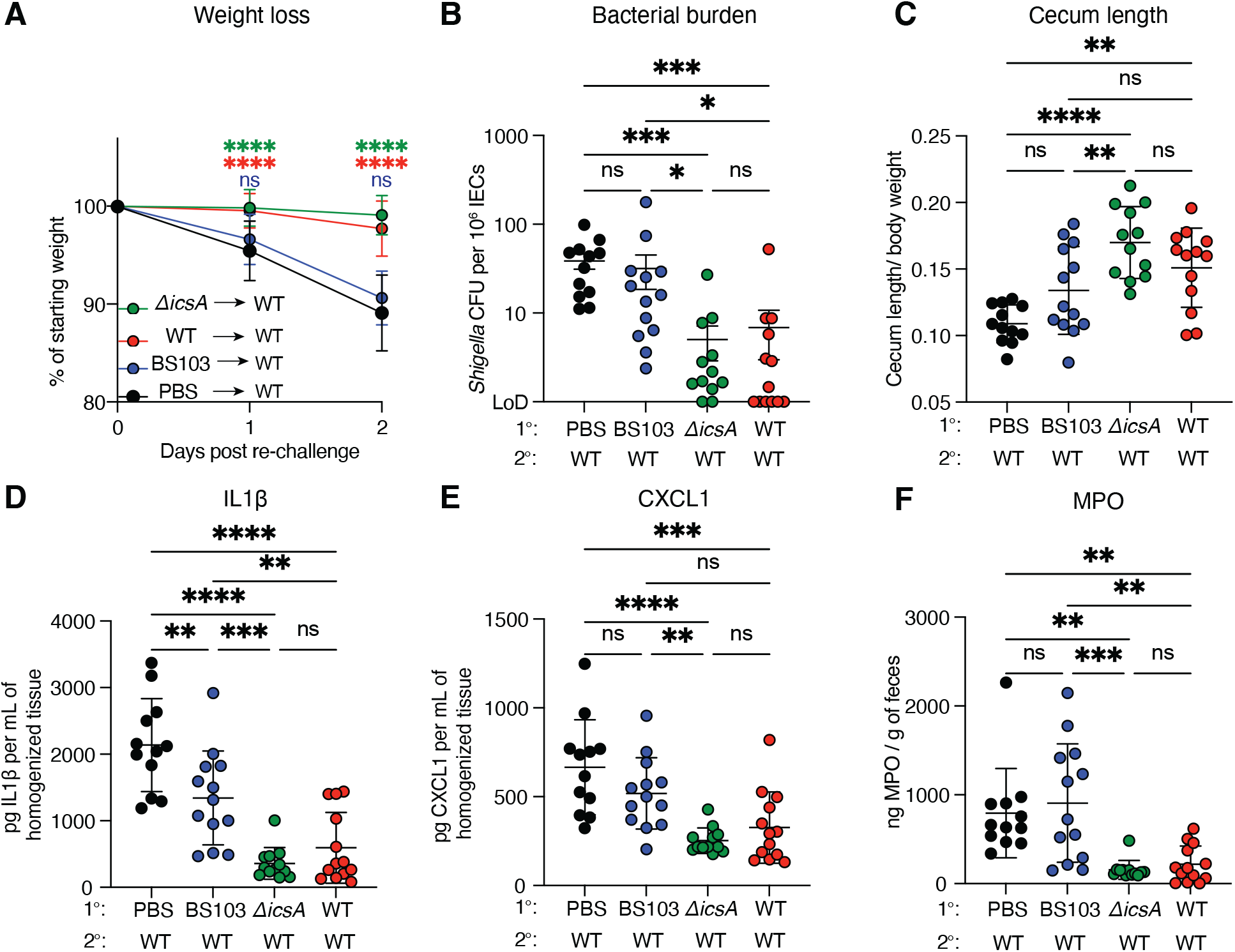
Virulence plasmid is necessary for protection against re-infection. (**A-F**) Mice were infected with WT *Shigella*, BS103 or D*icsA*. Mice re-challenged with WT *Shigella*. (**A**) Mouse weights from 0 to 2 days post re-challenge. Each symbol represents the mean of mice of the indicated infection group. (**B**) *Shigella* CFUs per million intestinal epithelial cells upon re-infection. (**C**) Cecum length normalized to the mouse weight before infection. (**D** and **E**) IL 1 and CXCL1 levels measured by ELISA from homogenized colon and cecum tissue. (**F**) MPO levels measured by ELISA from homogenized feces. Each symbol represents an individual mouse (**B-F**). Data are pooled from two independent experiments. Mean ±SD is shown in (**A, C-F**). Mean ±SEM is shown in (**B**). Statistical significance was calculated by two-way ANOVA with **Tukey’s** multiple comparison test (**A**), Kruskal-Wallis test (**B**) and ordinary one-way ANOVA (**C-F**). *p<0.05, **p<0.01, ***p<0.001, ****p<0.0001, ns = not significant (p>0.05).

**Supplementary Figure 4.**
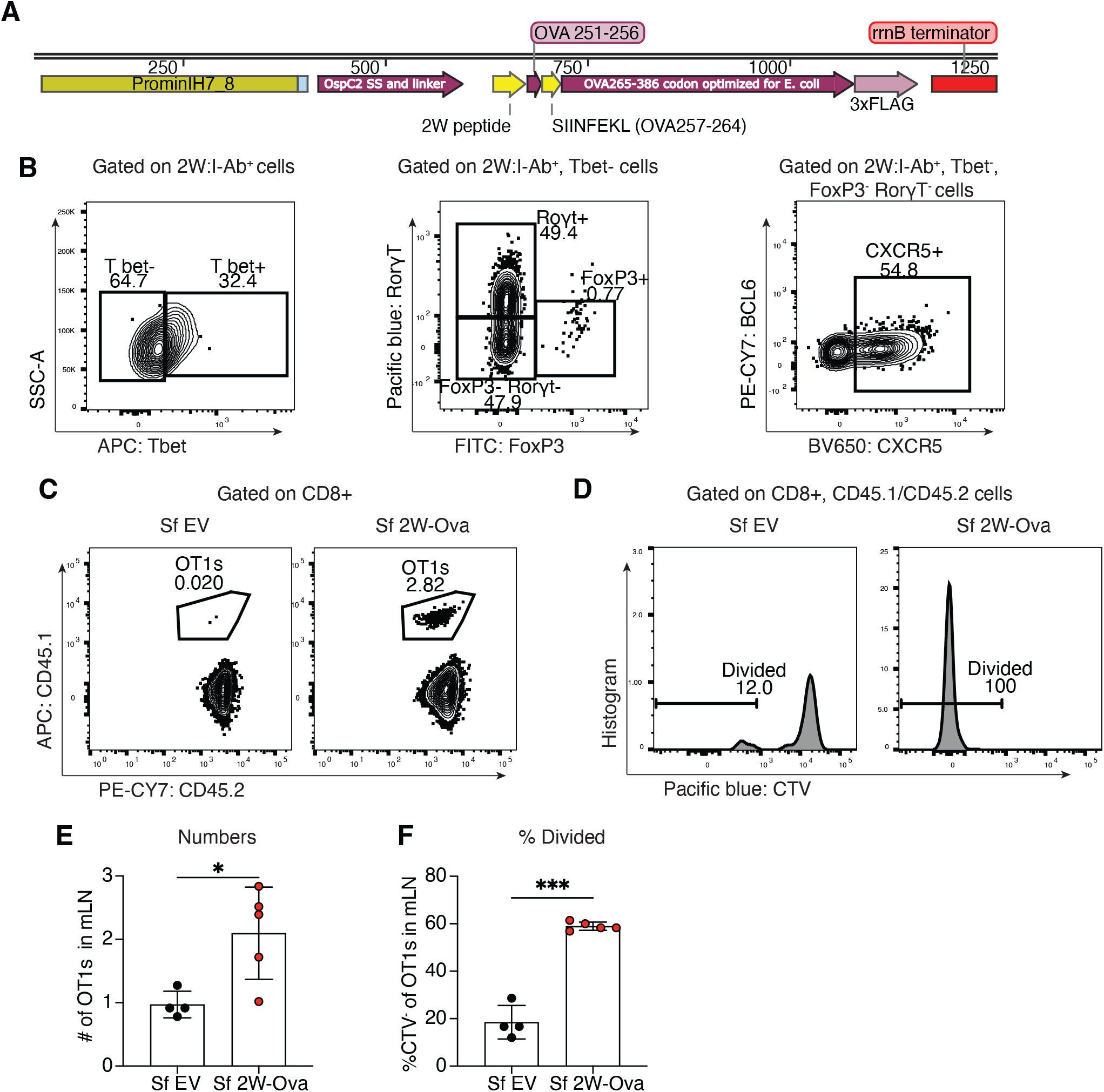
Primary *Shigella* infection induces CD4^+^ and CD8^+^ T cell expansion and activation. (**A**) Construct of FLAG-tagged 2W and Ova fused to the secretion signal of the OspC2 effector used to generate Sf 2W-Ova. (**B**) Mice were infected with *Sf* 2W-Ova, *Sf* EV or intraperitoneally injected with 2W peptide and polyI:C (IP) as a positive control. Splenocytes and mesenteric lymphocytes (mLN) were obtained and magnetically enriched for 2W^+^ CD4^+^ T cells using a tetramer, and then analyzed by flow cytometry, 7 days post infection. Representative flow plots for transcriptional factor staining. OT1 cells (CD45.1/CD45.2) from a transgenic mouse were retro-orbitally transferred into recipient mice (CD45.2). Mice were orally challenged with *Sf* 2W-Ova or *Sf* EV, and CD8^+^ T cells analyzed by flow cytometry. (**C**) Representative flow plots for transferred OT1 cells in recipient mice 7 days post infection. (**D**) Representative flow plots for proliferation of OT1 cells and (**E, F**) Quantification.

**Supplementary Figure 5.**
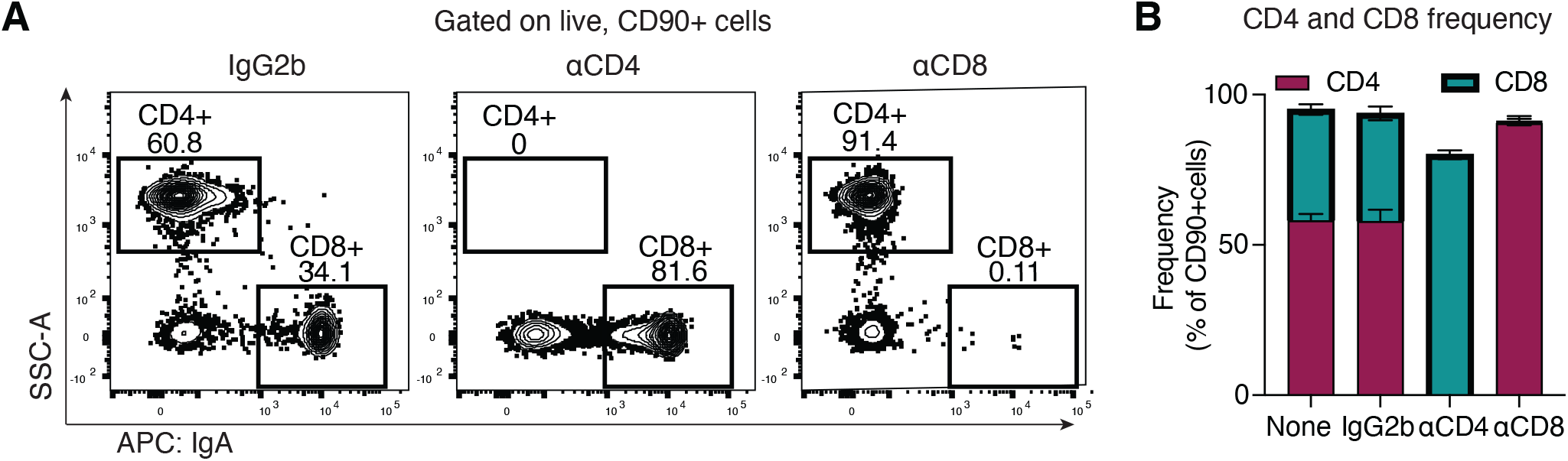
CD4^+^ T cells and CD8^+^ T cells are depleted. Mice were IP injected with anti-CD4 (αCD4) or anti-CD8 (αCD8) depleting antibodies 3 days prior to primary infection with WT *Shigella*, and every 2 days post-infection for 2 weeks. Control mice received an IgG2b isotype control antibody. Mice were bled 13 days post infection and analyzed for CD4^+^ and CD8^+^ T cell frequencies by flow cytometry. (**A**) Representative flow plots. (**B**) Quantification.

**Supplementary Figure 6.**
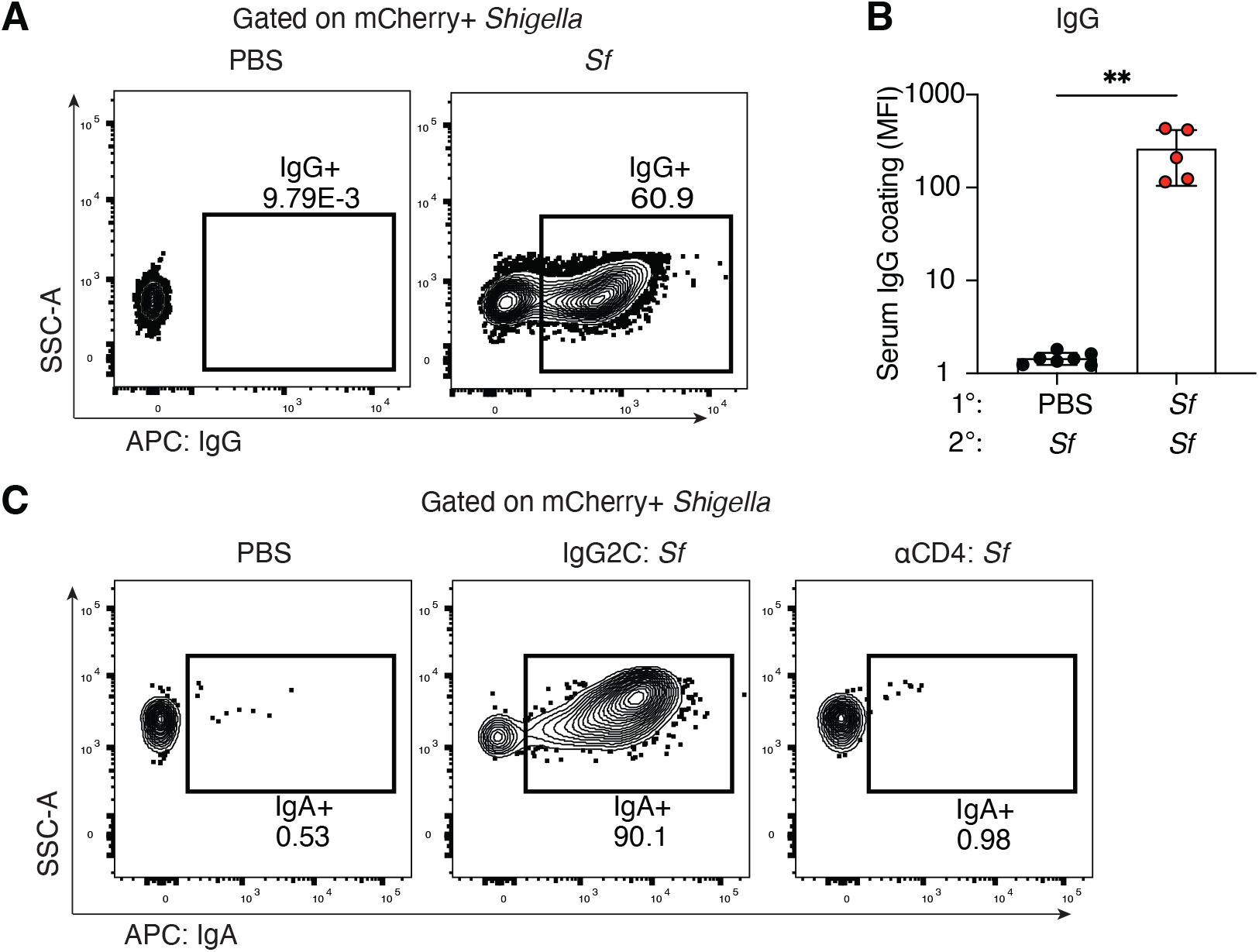
CD4-dependent antibody responses against *Shigella*. (**A-B**) Mice were infected as outlined in Fig. 1 with WT *Shigella* or PBS. Mice were re-challenged with WT *Shigella* expressing mCherry. Blood was obtained from mice and analyzed for *Shigella*-specific IgG by flow cytometry. (**A**) Representative flow plots for *Shigella*-specific IgG in serum. (**B**) Quantification (**C**) Mice were intraperitoneally administered with anti-CD4 depleting antibody 3 days prior to primary infection with WT *Shigella* and every 2 days post infection for 2 weeks. Control mice received an IgG2b isotype control antibody. Mice were re-challenged with WT *Shigella*. Representative flow plots for IgA coating following CD4^+^ T cell depletion. Statistical significance was calculated by Mann-Whitney test (**B**)*p<0.05, **p<0.01, ***p<0.001, ****p<0.0001, ns = not significant (p>0.05).

**Supplementary Figure 7.**
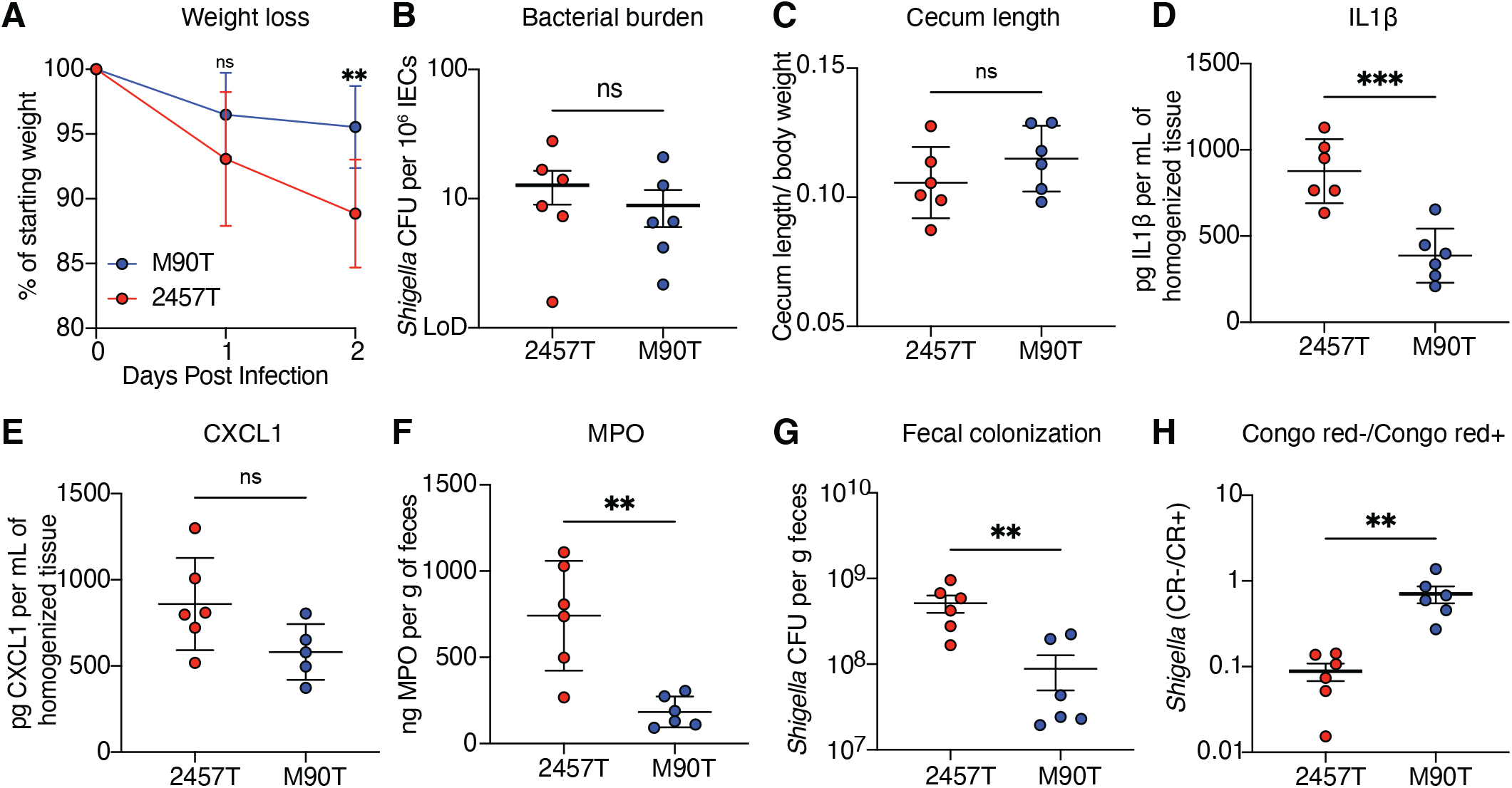
M90T exhibits lower virulence as compared to 2457T. (**A-H**) Mice were infected with *S. flexneri* M90T or 2457T and assessed for disease severity 2 days post infection (**A**) Mouse weights from 0 to 2 days post infection. Each symbol represents the mean of mice of the indicated infection group. (**B**) *Shigella* CFUs per million intestinal epithelial cells upon primary infection. (**C**) Cecum length normalized to the mouse weight before infection. (**D** and **E**) IL 1b and CXCL1 levels measured by ELISA from homogenized colon and cecum tissue. (**F**) MPO levels measured by ELISA from homogenized feces. (**G**) *Shigella* CFU per gram of feces. (**H**) Ratio of Congo red negative to positive *Shigella*. Each symbol represents an individual mouse (**B-H**). Data are representative of two independent experiments. Mean ±SD is shown in (A, C-F). Mean ±SEM is shown in (B, G and H). Statistical significance was calculated by two-way ANOVA with Tukey’s multiple comparison test (A), Kruskal-Wallis test (B, G and H) and ordinary one-way ANOVA (C-F). *p<0.05, **p<0.01, ***p<0.001, ****p<0.0001, ns = not significant (p>0.05).

**Supplementary figure 8.**
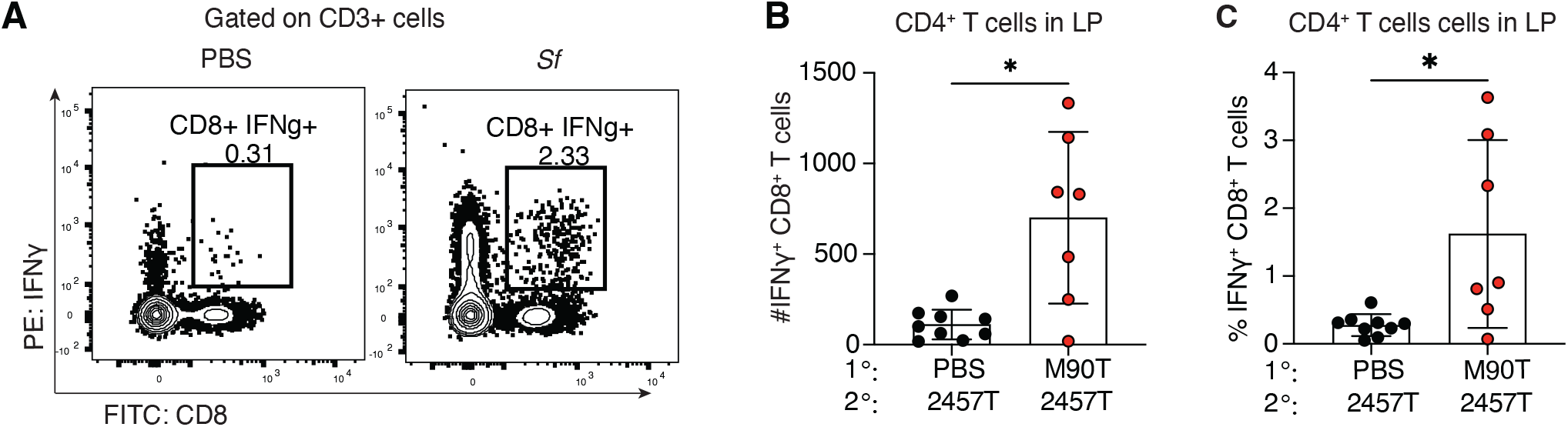
CD8^+^ T cells, although not required for protection on re-challenge, produce IFN-*γ*. Mice were intravenously injected with brefeldin A 1 day post re-infection with 2457T and IFN*γ*-producing CD8^+^ T cells in the lamina propria were quantified. (**A**) Representative flow plots of IFN*γ*-producing CD8^+^ T cells in the lamina propria (LP). (**B**) Number and (**C**) frequency of IFN*γ*-producing CD8^+^ T cells in the LP.

**Supplementary Table 1.** Antibodies.

| Target | Designation | Source | Cat | Dilution |
| --- | --- | --- | --- | --- |
| Surface | Ghost Dye Red™ 780 | Tonbo | 13-0865-T100 | 1/1000 |
| Surface | Anti-Mouse CD11c APC-eFluor™ 780 | eBioscience |  | 1/400 |
| Surface | Anti-Mouse CD11b (M1/70), APC-eFluor™ 780 | eBioscience | 47-0112-82 | 1/400 |
| Surface | Anti-Mouse CD45R (B220), APC-eFluor™ 780 | eBioscience | 47-0460-82 | 1/400 |
| Surface | Anti-Mouse CD4 BV786 | BD Horizon | 563727 | 1/200 |
| Surface | Anti-Mouse CD90.2 BUV395 | BD Horizon | 566557 | 1/1000 |
| Surface | Anti-mouse/human CD44 Antibody Alexa Fluor® 700 | Biolegend | 103026 | 1/200 |
| Intracellular | APC anti-T-bet Antibody | Biolegend | 644813 | 1/100 |
| Intracellular | BV421 Mouse Anti-Mouse RORyt | BD Horizon | 562894 | 1/100 |
| Intracellular | Anti-mouse FOXP3 Antibody Alexa Fluor® 488 | Biolegend | 126406 | 1/100 |
| Intracellular | Anti-Mouse Bcl-6 PE-Cy™7 | BD Biosciences | 563582 | 1/100 |
| Surface | Anti-Mouse CD62L (BV786) | eBioscience |  | 1/200 |
| Surface | Anti-mouse CD45.1 APC | eBioscience | 17-0453-81 | 1/200 |
| Surface | Anti-mouse CD45.2 PE/Cy7 | BioLegend | 109830 | 1/200 |
| Surface | Anti-mouse CD8a Brilliant Violet 650 | BioLegend | 100742 | 1/200 |
| Surface | Anti-mouse CD44 BB515 | BD | 564587 | 1/200 |
| Surface | Anti-mouse CD8 FITC | BioLegend | 100706 | 1/200 |
| Surface | Anti mouse CD3 APC | eBioscience | 17-0031-81 | 1/200 |
| Surface | Anti-mouse CD45 BV421 | BioLegend | 103134 | 1/200 |
| Surface | Anti-mouse NK1.1 BV650 | BioLegend | 108736 | 1/200 |
| Intracellular | Anti-mouse IFN-γ PE | BD Pharmingen | 554412 | 1/100 |

